# Continuous multiplexed population representations of task context in the mouse primary visual cortex

**DOI:** 10.1101/2021.04.20.440666

**Authors:** Márton Albert Hajnal, Duy Tran, Michael Einstein, Mauricio Vallejo Martelo, Karen Safaryan, Pierre-Olivier Polack, Peyman Golshani, Gergő Orbán

## Abstract

Primary visual cortex (V1) neurons integrate motor and multisensory information with visual inputs during sensory processing. However, whether V1 neurons also integrate and encode higher-order cognitive variables is less understood. We trained mice to perform a context-dependent cross-modal decision task where the interpretation of identical audio-visual stimuli depends on task context. We performed silicon probe population recordings of neuronal activity in V1 during task performance and showed that task context (whether the animal should base its decision on visual or auditory stimuli) can be decoded during both intertrial intervals and stimulus presentations. Context and visual stimuli were represented in overlapping populations but were orthogonal in the population activity space. Context representation was not static but displayed distinctive dynamics upon stimulus onset and offset. Thus, activity patterns in V1 independently represent visual stimuli and cognitive variables relevant to task execution.

## Introduction

To survive and thrive, organisms need to respond in distinct ways to identical stimuli, depending on the wider behavioral context. How the brain represents and maintains contextual variables during decision making is poorly understood. Specifically, it is not known whether contextual information is solely maintained in higher order brain regions such as frontal or parietal regions or if this information is fed back to primary sensory cortices to adapt perception to context. Task context can be considered as the variables that determine the contingencies between stimulus and rewards, but these variables may not necessarily be directly observed; rather the animal may infer the value of these variables and store them internally. Devising optimal actions might therefore require the confrontation of current sensory observations with the internally maintained latent state. Recent studies have demonstrated that non-sensory, task-related variables are represented in the primary visual cortex of rodents (Keller, Bonhoeffer & Hübener, 2012; Niell & Stryker, 2010; Poort et al., 2015; Shuler & Bear, 2006) and population responses undergo task-related transformations when performing a visual task (Goltstein et al., 2013; Meijer et al., 2017). If such task-related transformations occur in the visual cortex then it is critical to understand how the visual cortex can efficiently contribute to performing multiple tasks. Specifically, it is important to determine whether latent states corresponding to internal expectations about the actual task context are represented in the visual cortex.

Structured activity is present in the visual cortex and in particular in V1 even in the absence of visual stimuli (Fiser, Chiu & Weliky, 2004). Previous studies have suggested that the statistical structure of neural activity in the absence of stimulus may relate to the subject’s prior expectations of future sensory inputs (Berkes et al., 2011) and maintenance of such a prior contributes to the interpretation of sensory experiences (Chater, Tenenbaum & Yuille, 2006; Kok et al., 2013). Similar to environmental features, immediate task context is also inferred from and affects the interpretation of sensory stimuli. Yet, its presence in off-stimulus activity has not been explored. Finally, off-stimulus activity has recently been reported to reflect movement-related variables (Stringer et al., 2019a). This movement related activity was shown to reside in a subspace that is orthogonal to that occupied by visually evoked activity, thus ensuring that processing of visual information remains relatively intact by ongoing movements. It remains an open question, however, if a context variable which could affect processing of visual stimuli is also represented in an independent fashion.

To investigate these questions, we designed a paradigm where mice were required to switch between two tasks during a single recording session. During both tasks, mice were presented with the same visual and auditory stimuli. The two tasks differed in the stimulus modality (audio or visual) on which the animals were required to base their decisions. We performed multi-channel electrophysiological recordings in V1 of mice during task performance. Critically, in our paradigm, identical stimuli lead to different behavioral outcomes depending on context. Hence, the latent variable related to the context (whether the animal based its choices on visual or auditory input) could be identified in the neuronal population activity. We addressed two questions: First, we investigated whether the latent variable ‘task context’ is represented during on-stimulus activity and how this representation affects the representation of the visual stimulus. Second, we analyzed neuronal activity during the inter-trial interval to determine if task context is represented during off-stimulus activity and whether this representation of task context changes during visual stimulation.

We found that context can be decoded with high accuracy from V1 population activity both during visual stimulus presentation and inter-trial intervals. This context representation was orthogonal to the representation of stimulus identity. Our analysis showed that despite consistent representation of context during the trials and in intertrial intervals, the representation was not static but displayed distinctive dynamics upon stimulus onset and offset. In summary, activity patterns in V1 independently represent visual stimuli and cognitive variables relevant to task execution.

## Results

### Non-visual variable related activity in V1

We trained mice on a cross-modal audiovisual go/no-go task (Fig. 1A). The task consisted of four blocks: two unimodal sensory discrimination blocks where animals licked or refrained from licking either based on the orientation of the visual stimulus (45 degrees or 135 degrees) or the frequency of an auditory tone (5 kHz or 10 kHz) and two cross-modal discrimination blocks where animals were presented with both auditory and visual stimuli and made their decision to lick based on one modality while ignoring the other. The unimodal and cross-modal blocks were interleaved and during cross-modal blocks, animals learned to base their decisions on the sensory modality that was presented during the preceding unimodal block. Critically, during the cross-modal blocks, the two stimulus modalities were presented simultaneously and it was the task context that determined the relevance of a particular modality with respect to the choice that the animals had to make (Fig. 1A). Only licking or refraining from licking during the final second of the 3 second visual stimulus was considered as the animal’s decision. Misses and false alarms led to a timeout. The stimulus modality in the initial block was selected randomly for any given day and the modality of the third block, the transition block, was the other modality.

**Figure 1.**
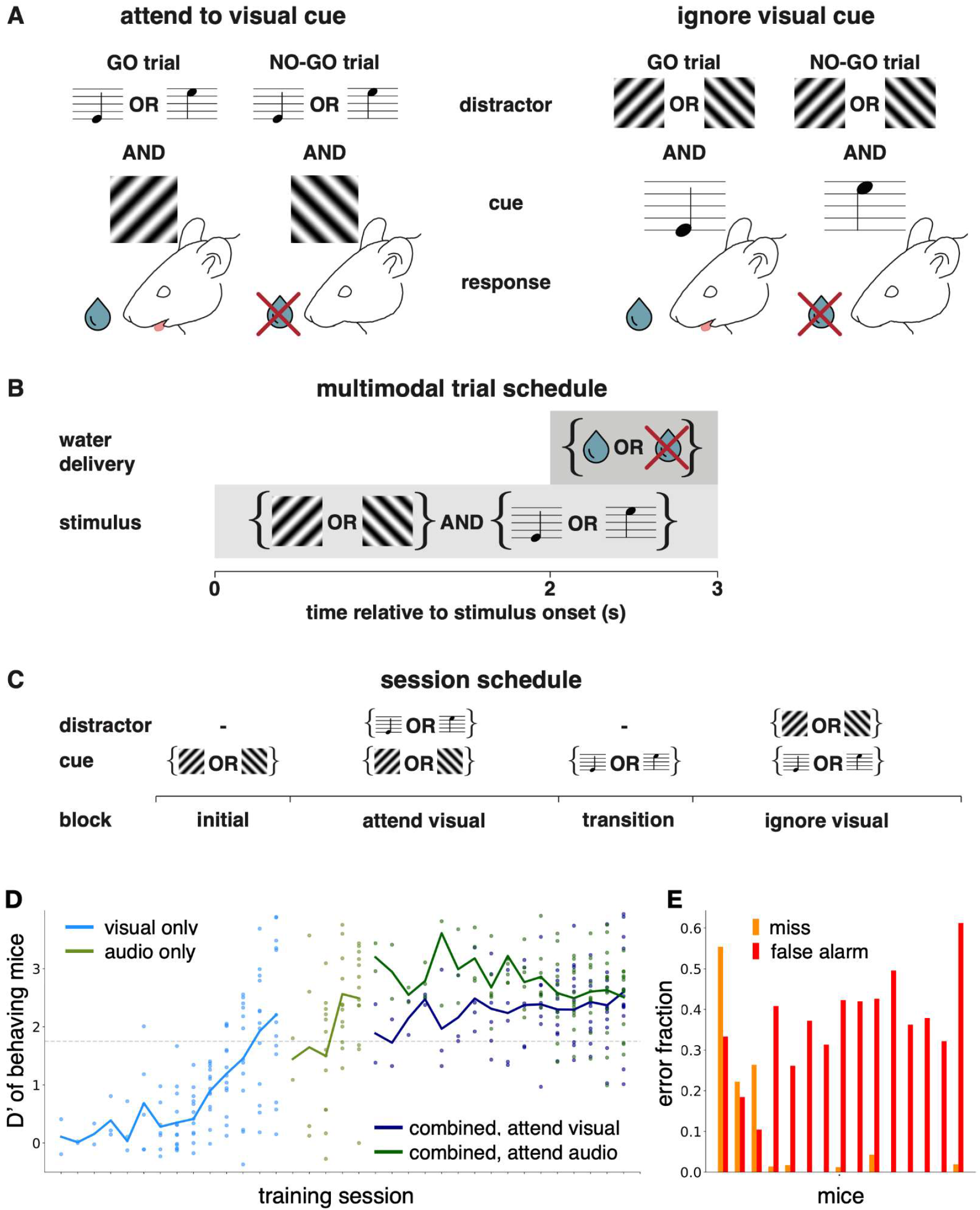
Behavioral paradigm. **A**, Stimulus and reward structure in the attend visual stimulus (*left*) and ignore visual stimulus (*right*) tasks. In both tasks either of two visual stimuli (45° or 135° moving grating stimuli) and either of two auditory stimuli (high or low pitch) is presented. In the ‘attend’ visual task one of the visual stimuli was designated as a go signal where licking was rewarded with water. Licking for the no-go signal was punished by a short time-out. The identity of the auditory stimulus was irrelevant. In contrast, in the ‘ignore’ visual task the visual stimulus identity was irrelevant and mice were required to base their judgement on the identity of the auditory stimulus. **B**, Trial structure. Stimuli were presented for three seconds, and a three seconds rest period followed (time-out extended the delay to ten seconds). Water was available after two seconds from start. **C**, Structure of an experimental session. Initially a block of trials was presented where a single stimulus modality was present. The stimulus modality used in these trials cued the type of multimodal task in the subsequent block of trials: the modality relevant for decisions in the multimodal trial was identical to the modality of stimulus used in the preceding block. After the first multimodal task comes the transition block. In this block, the animals needed to transition to the other task context, where the previous distractor became the relevant stimulus modality. Similar to the initial block, animals were cued with the new relevant modality stimulus in this unimodal cueing transition block. The order of relevant stimulus modality between the two initial parts was randomized across animals. **D**, D-prime values of 13 animals representative of the whole mice population partaking in the experiments, as they first learn the visual discrimination task (light blue), then the auditory discrimination task (light green), and then perform the cross-modal attention task in the attend visual condition (dark blue), and attend auditory (dark green) condition, on average well above the target D’=1.75 expert level (dashed horizontal line). Each dot represents the averaged D-prime of one animal in a given training session. Solid lines are averages over D-primes of mice. Most mice were trained for fewer sessions than the full width of the three session types. All mice aligned to the last training session for the session type, hence fewer dots in the first few training session data points. **E**, Rate of misses (orange) and false alarms (red) in the 15 animals during the recording session.

We trained 15 animals to reliably perform the context-dependent decision making task (Fig. 1D). Mice were first trained on the visual discrimination task over 7 training sessions on average (Fig. 1D, green trace, see Methods for details), the auditory discrimination task over the subsequent 3 training sessions (Fig. 1D, gray trace), and then attend-to-visual and attend-to-auditory over 4 training sessions (Fig. 1C, red and blue traces, respectively). Animals performed 300-500 trials daily. Performance was measured using the d’ statistic which compares the standard deviation from chance performance during lick and no-lick trials (chance d’=0). Animals were considered experts if their sessions averaged d’ > 1.75 (probability of chance behavior < 0.1%). All animals showed average d’ values above the expert threshold of 1.75 in all behavioral tasks, demonstrating that they could be trained to learn the multimodal attention task reliably (Fig. 1D).

Neural activity was recorded from all V1 layers on 128 channels (two 64 channel shanks) with extracellular silicon probes (Du et al., 2011). Spiking activity of separated units was obtained by applying kilosort2 (Pachitariu et al., 2016) followed by strict manual curating. Importantly, task context (whether the animal is basing its decisions on the visual or auditory stimulus) is characterized by slow dynamics; this necessitates specific attention to effects that might correlate with the context variable. In particular, electrode drift could also potentially introduce long time-scale noise in the recorded activity which confounds the analysis of context representation. To prevent this, we relied on a conservative spike sorting approach. Units were tested individually for drift by assessing the constancy of signal to noise ratios on amplitudes, feature projections, and spiking frequency across the recording session (see Methods and Supplementary Fig. 1). Units that showed signatures of drift even on a short part of the session were discarded from further analysis, yielding 276 units across all mice.

During task performance, visual stimulus presentation to the monocular contralateral visual field induced substantial variance in the responses of recorded V1 neurons (mean firing rate changes from 8.93 ± 0.70 to 10.17 ± 0.77 Hz from baseline to stimulus presentation, trial to trial variance on stimulus, mean and s.e.m. over neurons: 29.19 ± 4.05 Hz^2^, number of neurons 276 from 15 mice) including neurons whose firing rate was increased (7.96 ± 0.66 to 11.34 ± 0.80 Hz and trial to trial variance on stimulus: 33.24 ± 3.50 Hz^2^, 171 neurons) or decreased (10.51 ± 0.76 to 8.25 +/- 0.70 Hz and trial to trial variance on stimulus: 22.59 ± 4.80 Hz^2^, 105 neurons), respectively (Fig. 2B). Mean waveform trough-to-peak time and amplitude clustering revealed 101 narrow spiking and 175 broad spiking (putative inhibitory and excitatory) units, with firing rate change to stimulus from 12.85 ± 1.62 to 15.04 ± 1.73 Hz and 6.67 ± 0.53 to 7.35 ± 0.58 Hz respectively. Visual stimulus identity was not the only task variable which induced variance in neural responses during visual stimulation. In fact, individual neurons showed sensitivity to visual stimulus identity, auditory stimulus identity, or the choice the animal was about to make (Fig. 2B). In addition, we also found neurons that were influenced by the decision making context, i.e. the modality the animal is required to base its choices on (Fig. 2B). Individual neurons were not selective to one specific task variable, rather varying levels of mixed selectivity could readily be observed (note differential selectivity to multiple variables on Fig. 2B).

**Figure 2.**
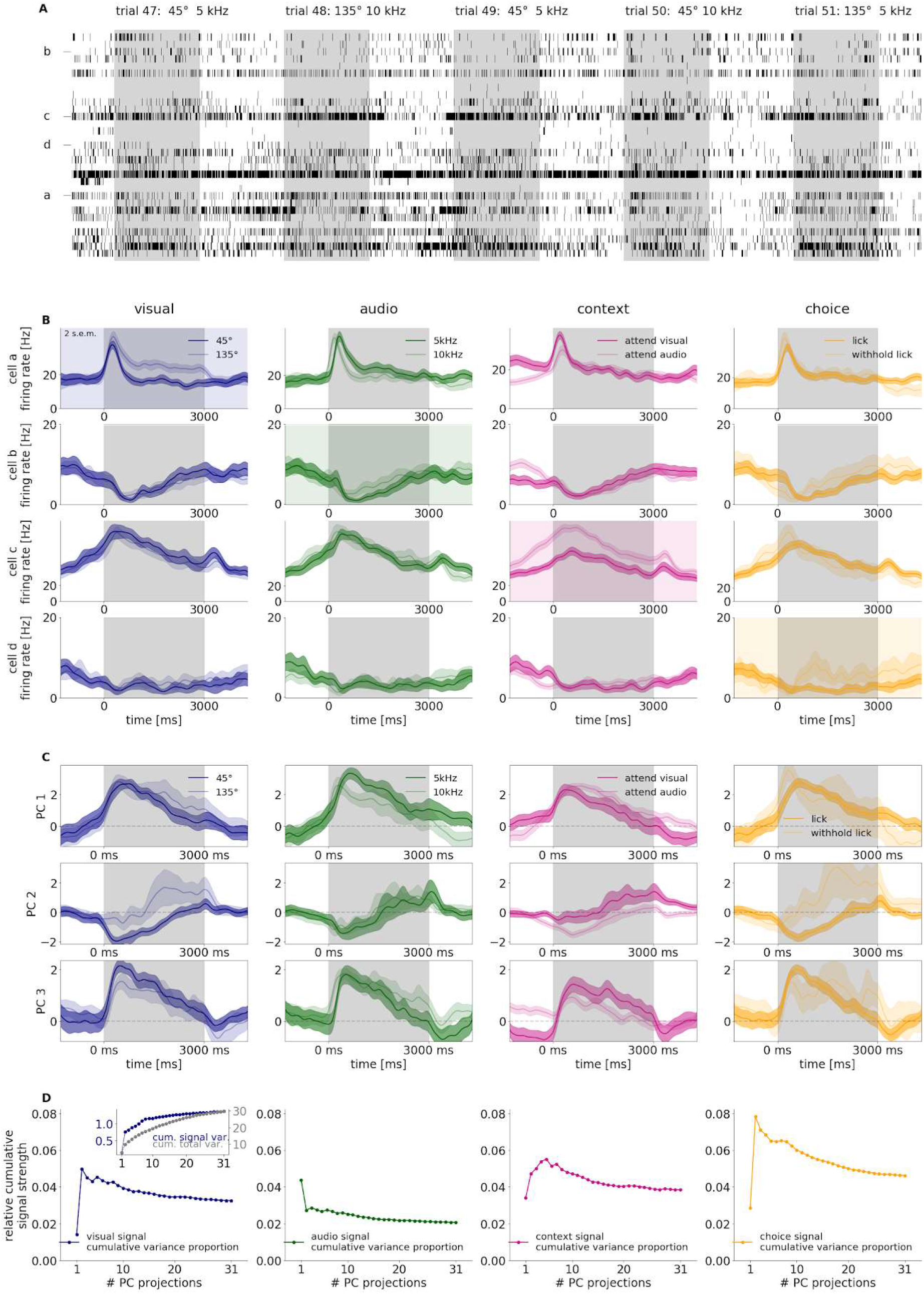
Population recording from the V1 of an example mouse. **A**, Spike raster for the recorded population of 31 neurons in five subsequent multimodal trials. Stimulus presentation windows are marked with *grey background*, while activity during the intertrial interval is marked by a *white background*. Letter labels indicate the neurons analyzed on panel **B**. **B**, Sensitivity of four example neurons (rows) to different task-relevant variables (columns). Across-trial average firing rates are calculated for two values for the particular variable (*dark and light lines, shadings* around lines denote 2 s.e.m.): 45° and 135° angle for visual stimulus, low and high pitch stimuli for auditory stimuli, attend visual and ignore visual task for context, and lick and withhold lick for choice. **C**, Task-relevant activity identified in population-wide analysis of the responses with Principal Component Analysis (PCA) performed on population responses in the full stimulus presentation period. Time course of population activity projected on the first three principal components (*three rows*) averaged across different values of the four task-relevant variables (*four columns*). Colors are matched with those on panel **B**. **D**, Relative signal variance of population activity associated to the state of the four different task-relevant variables (*four columns*). Variance is normalized by the total variance of the population responses. Relative variances are shown cumulatively for activities projected on a growing subset of PCs. Inset on a separate vertical scale shows signal variance, the numerator and total variance, the denominator of the relative variance shown in the main panel.

Selectivity of individual neurons to different task variables indicates that all relevant variables are represented in the population but it remains unclear how dominant the activity related to these task variables is in relation to the total population activity. To capture the relative contribution of the variance related to different task variables to the total variance in population activity, we used an unsupervised method, principal component analysis (PCA), to identify orthogonal dimensions (principal components, PCs) in the population activity in which activity displays large across-trial variance and to rank these dimensions according to the magnitude of variance associated with them. Activity along individual PCs showed that task variables can be readily identified in the population activity, as revealed by calculating mean population activity along individual PCs for different conditions of task variables (Fig. 2C). To assess the relative contribution of the task variables to population activity, we calculated the variance explained by individual task variables (termed signal variance) for each PC. For this, we first assessed the signal variance in individual PCs and then the cumulative signal variance was established as the sum of variance explained by the specific task variable in a subset of PCs (Fig. 2D, inset, see also Methods for details). Finally, we calculated the relative contribution of a specific task variable to the total variance in the population by calculating the ratio of cumulative signal variance and the total variance captured by the same subset of PCs. This method is similar to demixed PCA (Kobak et al., 2016), but differs in that our approach treats task variables separately. The advantage of this analysis technique is that it does not make assumptions about task variable dependencies (see also Methods). The relative cumulative signal variances peaked for low dimensional PC subspaces, indicating that all task variables contributed to the dominant modulation of population activity (Fig. 2D). The relative cumulative signal variance gradually decreased when the rank of the PC dimension increased and summed to a few percent for the whole analyzed population. In summary, population activity in V1 is influenced by all relevant task variables and these task variables contribute to the leading directions of variance.

### Orthogonal representation for visual stimuli and task context

The above analyses highlighted that the task context is represented in the population activity of V1 neurons. To further understand how the visual cortex differentially represents identical stimuli when these stimuli indicate different behavioral outcomes, we constructed linear decoders (Fig. 3A) for visual stimulus identity (Fig. 3B) and context (Fig. 3E). A linear decoder identifies a hyperplane in the linear activity space of the neuronal population which can most efficiently separate the population responses evoked by two particular values of a relevant variable (Fig. 3A). We could then determine a decision vector (DV) orthogonal to the hyperplane, that identified the direction along which the neuronal activity changed most when the value of the decoded variable changed. A time-dependent measure of the strength of the representation was obtained by fitting separate decoders for 50 ms of neural data recorded in individual animals (Fig. 3B,E, for details see Methods). To distinguish decodability from chance we compared decoder accuracies to randomized label decoders (see Methods), and are highlighted as grey dashed lines on relevant figure panels. Both visual stimulus content and task context can be decoded throughout the stimulus presentation period, and this is consistent across animals (Fig 3C,F, mean visual accuracy 0.68 ± 0.04, 2 s.e.m. of visual accuracy 0.03 ± 0.00, mean context accuracy 0.65 ± 0.03, 2 s.e.m. 0.03 ± 0.00, number of animals: 15). As parallel representation of visual stimulus identity and task context can be achieved by separate subpopulations of neurons selective for one variable or the other, we assessed the overlap between these subpopulations by comparing the contribution of individual neurons to the two linear decoders (Fig. 3D, G). Decoder weights for visual stimulus were well distributed across the analyzed neuronal population and showed strong overlap with the weights of the task context decoder. This finding implied that neurons presented a mixed selectivity for stimulus and context. As GABAergic interneuron classes have been reported to mediate top-down signals in sensory areas (Zhang et al., 2014), it was therefore important to test if the context signal we recorded was not solely a result of activity in interneuron populations. Putative principal cells and interneurons were distinguished based on their spiking waveforms (see Methods). We tested if the context representation is affected by constraining our analysis to broad spiking, putative principal, cells. We only analyzed mice in which the size of the available neuronal population was larger than 30 and the number of broad spiking neurons larger than 15. We found that accuracy of decoding context from broad spiking neurons was close to that from all neurons. (Supplementary Fig. 2) indicating that context is jointly represented in excitatory and inhibitory neuron populations.

**Figure 3.**
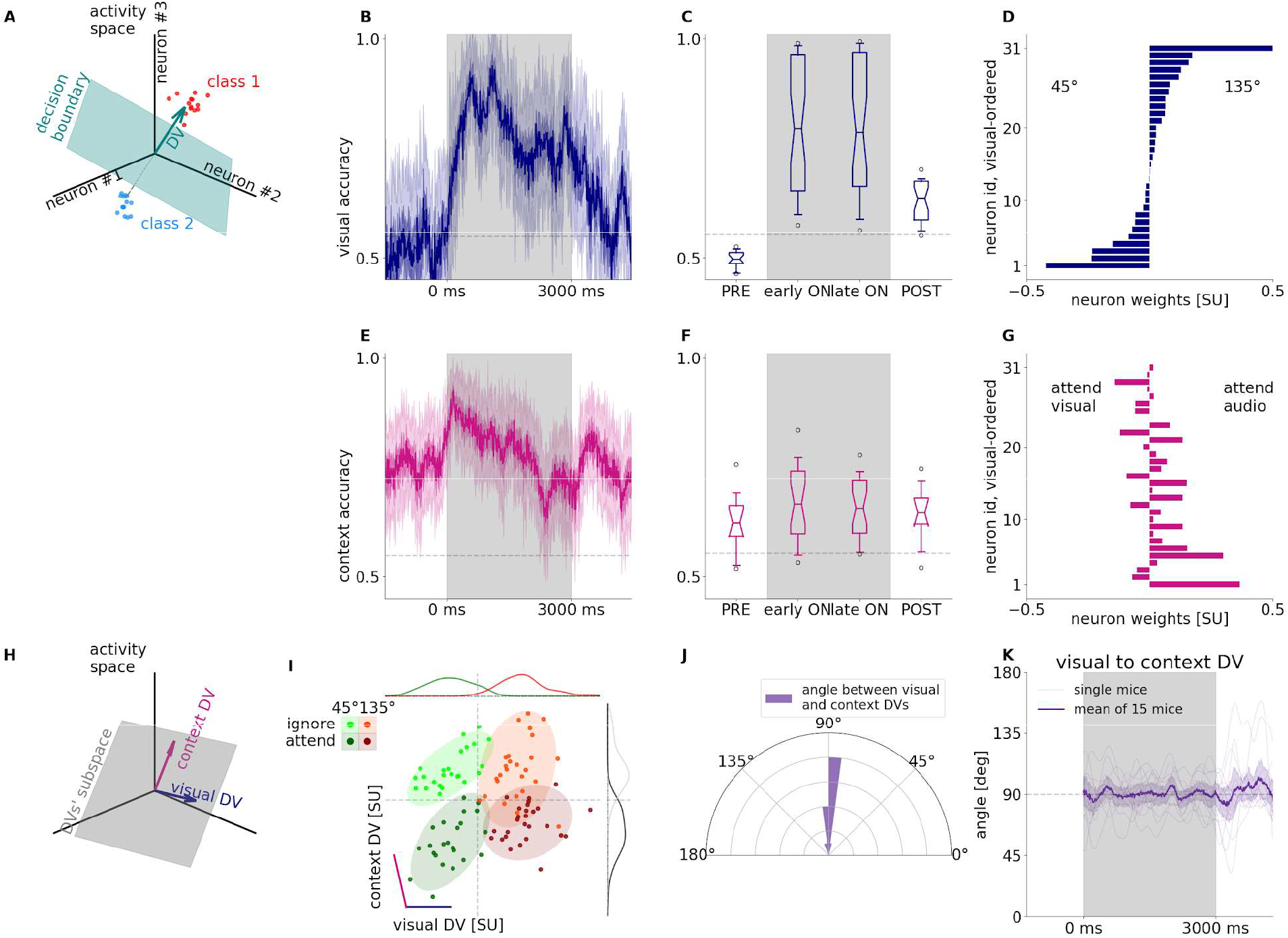
Representation of task context in population activity. **A**, Illustration demonstrating linear decoding from population activity. Linear decoding analysis discovers a decision boundary (green surface) in the activity space which separates responses measured in individual trials (*dots*) in two conditions (red and *blue*). This linear subspace is a two-dimensional plane when analyzing the responses of three neurons. The decision boundary can be uniquely characterized by the decision boundary normal vector (decision vector, DV, *green arrow*), a vector perpendicular to the boundary surface. **B**, Decoder performance in 50 ms time windows at 10 ms sliding resolution for visual stimulus identity, prior to stimulus onset (PRE), during stimulus (ON), and after stimulus (POST) for an example animal. Gray shading marks the stimulus presentation period. Grey dashed line indicates 1 s.e.m. of randomized labels decoder accuracies around chance level. Note that firing rates are estimated by using convolution kernels with 100-ms characteristic width and therefore decoders contain information from future time points. This residual contribution results in a slight increase in visual decoding performance prior to stimulus onset. **C**, Average performance of visual decoder for all animals. Box and whiskers denote 25-75, and 2.5-97.5 percentiles respectively, midlines are the mean, notches are 95% confidence level error of the mean. Grey dashed line indicates 1 s.e.m. of randomized labels decoder accuracies around chance level averaged over animals. **D**, Contributions of individual neurons (decoder weights) arranged according to the magnitude of the weights. **E-G**, same as **B-D** but for decoding context from the population activity. Ordering of neurons on panel **G** is the same as that on panel **D**. **H**, Combination of multiple DV bases forms a new basis defining a higher dimensional subspace of task relevant population activity. **I**, Population responses projected on the DV subspace in individual trials (*dots*) in different task contexts (*dark* and *light*) and with different visual stimuli presented (red and *green*). Purple and blue lines denote the DBNV directions of context and visual decoders, respectively. Histograms show population responses projected on orthogonal components of single DVs. Trials come from multimodal trials in an example animal. **J**, Histogram of the angle between context and visual DVs across animals using average activity in the first 1.5 s of stimulus presentation. **K**, Time course of the angle between the visual and context decoders throughout the trial.

A context representation that is not independent from visual content representation could yield correlations that are detrimental to the accuracy of the visual stimulus identification. We therefore investigated the relationship between the representations of the two task-relevant variables by analyzing the properties of the decoders trained to distinguish context and visual content. Using our Multidecoder Subspace Analysis framework (MDSA, see also Methods), we constructed a trial-by-trial measure of population activity: population vectors constructed from spike counts of recorded neurons during the first second after stimulus onset were projected on a two-dimensional subspace defined by the two directions along which the population activity could be separated most for the two investigated variables, context and visual content (Fig. 3H). This two-dimensional subspace was a generalization of more traditional analyses which project activity vectors onto the one-dimensional subspace (i.e. line) defined by the DV of a single decoded variable as basis, by linearly combining two of such bases in the same population activity space. Population responses were distributed along the direction which most effectively distinguishes population responses in two conditions separately (Fig. 3I, marginals). The two-dimensional generalization of this concept used the plane defined by the DVs of two variables calculated for the first second after stimulus onset. (Fig. 3I). Population activity in individual trials (Fig. 3I, dots) revealed how the response related to the response variance corresponding to both of the variables. Responses in multimodal trials showed a striking pattern: population responses to different task contexts and different visual content formed distinct clusters, and variations across contexts and stimulus identities were close to orthogonal, highlighting that these variables were represented in independent subspaces albeit with overlapping populations (Fig. 3I). The relative independence of context and visual content representations is shown across animals: while the angle between the context and visual content decoders slightly varies, the distribution of decoder angles peaks close to 90° (Fig. 3J). Time-resolved analysis of the DV angle showed that orthogonality of visual content and task context is consistent across the stimulus presentation (Fig. 3K, see Methods for details). This finding is corroborated by decoding context from neural activity that is projected onto the 1 dimensional subspace of visual DV at each timepoint: Albeit precise orthogonality between visual and context related activity in single trials at every time point is not always achieved, generally decoding context from the visually relevant subspace yields close to chance accuracy (Supplementary Fig. 3).

### Context representation in the absence of stimulus

Task context is not independent across trials since context remains the same in a given block. As a consequence, we hypothesized that the neural representation associated with task context could not only be present during stimulus presentation but also during the intertrial interval. By training separate cross-validated linear decoders in a shifting time window, time-resolved linear decoding analysis demonstrated that decodability of context is persistent and can be decoded during pre-stimulus and post-stimulus activities with accuracies matching context decoding during stimulus presentation (Fig. 3B, pre-stimulus mean accuracy across animals 0.62 ± 0.03, during stimulus mean 0.66 ± 0.04, post-stimulus mean 0.65 +/- 0.03). We found significant variance in decoding accuracy between animals. However, decoding accuracy measured before and during stimulus presentation were strongly correlated (Fig. 4A, R=0.84, p=0.0001). By controlling for the number of units in the analysis, we confirmed that this dependency was not solely dependent on the number of neurons recorded (Supplementary Fig. 4).

**Figure 4.**
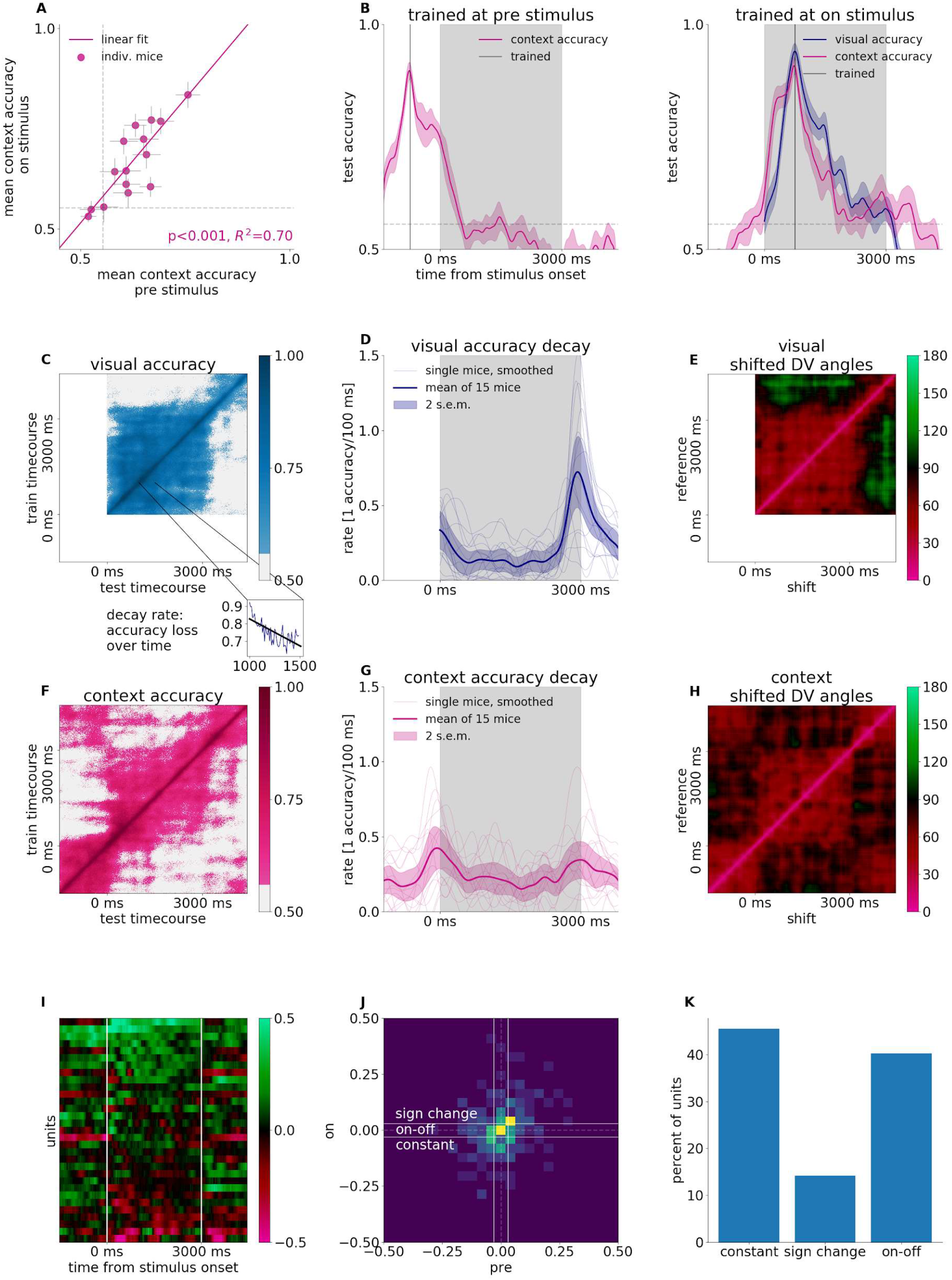
Context representation in the absence of stimulus. **A**, The relationship of context decoding performance in the intertrial interval (prior to stimulus presentation) to context decoding performance during stimulus presentation. Grey dashed line indicates 1 s.e.m. of randomized labels decoder accuracies around chance level. **B**, Accuracy of cross-testing visual (*blue*) and context (magenta) decoders at timepoints different from the trained timepoint (grey vertical line) at pre- (left) and on (*right*) stimulus, grey shaded area accounts for stimulus presentation. Grey dashed line randomized chance level as in **A**. **C**, Generalization capacity of the visual stimulus decoder across time. A decoder trained at different time windows during the trial (*vertical axis*) is tested at all the time windows during the trial (*horizontal axis*). Color scale shows decoder performance. Grey color indicates decoder performance under 1 s.e.m. of randomized labels decoder accuracies around chance level. *Inset*, a cross section of the main panel with accuracy color code transformed to vertical axis, shows the dependence of the decoder performance as a function of the delay of the test relative to the training. Black line shows a line fit characterizing the rate of decay. **D**, Smoothed rate of decay at each timepoint, linear fits on 500 ms forward as shown in the inset of **C** at 1000 ms. Thin lines show individual mice, thick line shows mean of 15 mice, shaded band shows 2 s.e.m. Grey shaded area accounts for stimulus presentation. **E**, Relative angle between the DBNVs of visual decoders trained at different time windows throughout the trial. **F-H**, Same as panels **C-E** but for the context variable. **I**, Context decoder coefficients (color code) at each timepoint for a representative animal. Neurons (vertical axis) ordered by mean coefficients during the on-stimulus period. **J**, 2D histogram (normalized color scale) of mean off-stimulus vs. on-stimulus decoder coefficients for (N=5, animals with number of units greater than 25). Units are categorized as opposite contributions in the two periods (*sign change*); contributing only to one of the periods (*on-off*); or contributing similarly to the two periods (constant). **K**, Distribution of units according to their differential contribution to on-stimulus and off-stimulus coding of context.

The consistency of context decodability during the pre-stimulus condition and on-stimulus conditions indicated that context representation was stable throughout the execution of the task irrespective of whether stimulus is presented or not. It was not clear, however, if the representations during pre-stimulus and on-stimulus activities were identical or undergo dynamical transformations. We investigated this question by constructing time-shifted decoders which assessed if the same DV could describe the changes in population activity with changes in the decoded variable at different time windows. Time shifted decoders were constructed by establishing the DV at a given time point during the trial and testing its efficiency at different time points during the trial (Fig. 4B). Time-shifted decoding showed that decoder performance slowly decayed as the difference between the time of training and testing grew both for the visual content decoder and for task context decoder. To understand how this decay is affected by switching between trial stages (pre-stimulus/on-stimulus/post-stimulus) we performed time-shifted decoding for all possible time shifts (Fig. 4C). This analysis yielded a two dimensional plot which showed the linear decoder performance for all possible training time windows and all possible delays. Time-shifted decoding of the visual content showed relatively consistent accuracy across the whole period of the stimulus presentation but performance dropped as the stimulus was switched off. We characterized the time course of the rate of decay by fitting a line to any particular time point in the time-shift-dependent decoding performance (Fig. 4C inset). As expected, the decay rate displayed a peak at the time when the stimulus was switched off (Fig. 4D). Consistency of the visual content related neural responses was also assessed by calculating the angle between DVs of visual decoders trained at different times of the trial (Fig. 4E): small angles corresponded to a more invariant code, while angles closer to 90 degrees corresponded to an independent code. DV angle dropped from close to 0 to about 45 degrees with small time difference between decoders but remained stable throughout the stimulus presentation.

Time-shifted analysis of task context spanned the pre-, on-, and post-stimulus periods (Fig. 4F). While time-shifted decoding resulted in above-chance performance during the stimulus presentation period, the decay of the time-shifted decoder was faster for task context than for the visual content (Fig. 4D and G, 15 mice, averaged forward decay rate in the steady state between 1000 and 2000 ms after stimulus onset, dependent t-test p=0.0491), indicating a gradual transformation that is faster for the context representation than the change measured for the visual representation. Importantly, during the pre-stimulus period the time-shifted decoding showed a relatively stable representation of task context (Fig. 4F, −1500 to −500 ms average decay rate of 0.19+/-0.04 accuracy loss /100 ms from 15 mice), but stimulus onset was characterized by an abrupt decay (Fig. 4G, 0.42+/-0.06 accuracy loss / 100 ms). This finding was corroborated by the analysis of DV angles, which showed small angles both within the pre-stimulus (45.3+/-3.7°, σ=14.3°) and within the on-stimulus periods (50.7+/-3.1°, σ=12.0°), but closer to orthogonal representation for task context across the border of pre- and on-stimulus activities (78.5+/-4.2°, σ=16.4°, all angle statistics from Fig. 4H, see Methods). In summary, while the task context is represented throughout the trial in V1, it undergoes a rapid transformation between stimulus onset and offset.

We investigated how much of this rapid transformation of representation at stimulus on and offset results from a possible change in subsets of active neurons encoding context. By visualizing decoder coefficients for each unit at each time point throughout the trial (Fig. 4I), qualitative differences between neurons emerge: There are units that have similar weights throughout the on-stimulus and off-stimulus periods, the coefficient of other units switches sign, indicating changes in response intensity in the two periods, while context coding units’ contribution diminishes in one of the periods (Fig. 4J, N=5 animals, having greater than 25 units available). The decoder coefficients of a subpopulation of neurons are smaller than a noise ceiling threshold (see Methods) indicating that these do not contribute significantly to context decoding. The proportion of neurons belonging to the qualitatively different groups are comparable to each other (Fig. 4K). In summary, the rapid change in context representation is a result of a population-wide transformation, encompassing both recruitment of new neurons and changes in activity in participating neurons, rather than an activity switch in distinct subpopulations (Fig. 4I).

### Independence of context and visual content related activities

Previous analysis highlighted that task context is represented both in the pre-stimulus and on-stimulus periods. Thus, in different task contexts the population activity differed between attend-visual and ignore-visual conditions at the time of the delivery of stimulus. This raises the possibility that different initial conditions of the population activity resulted in different population dynamics, or in other words: the activity pattern elicited by the same stimulus in two different task contexts differed. We analyzed this question by examining a subspace of the population activity in which the most prominent visual content and task context related activity occurred: the two-dimensional space outlined by the visual content DV and task context DV constructed by averaging DVs during the first second after stimulus onset. We calculated population trajectories in the two task contexts for the two different visual stimuli (Fig. 5A,B). Initial conditions of population trajectories showed consistent differences across contexts (filled circles) and the time course of the population trajectory during stimulus presentation showed markedly different patterns depending both on task context and stimulus content.

**Figure 5.**
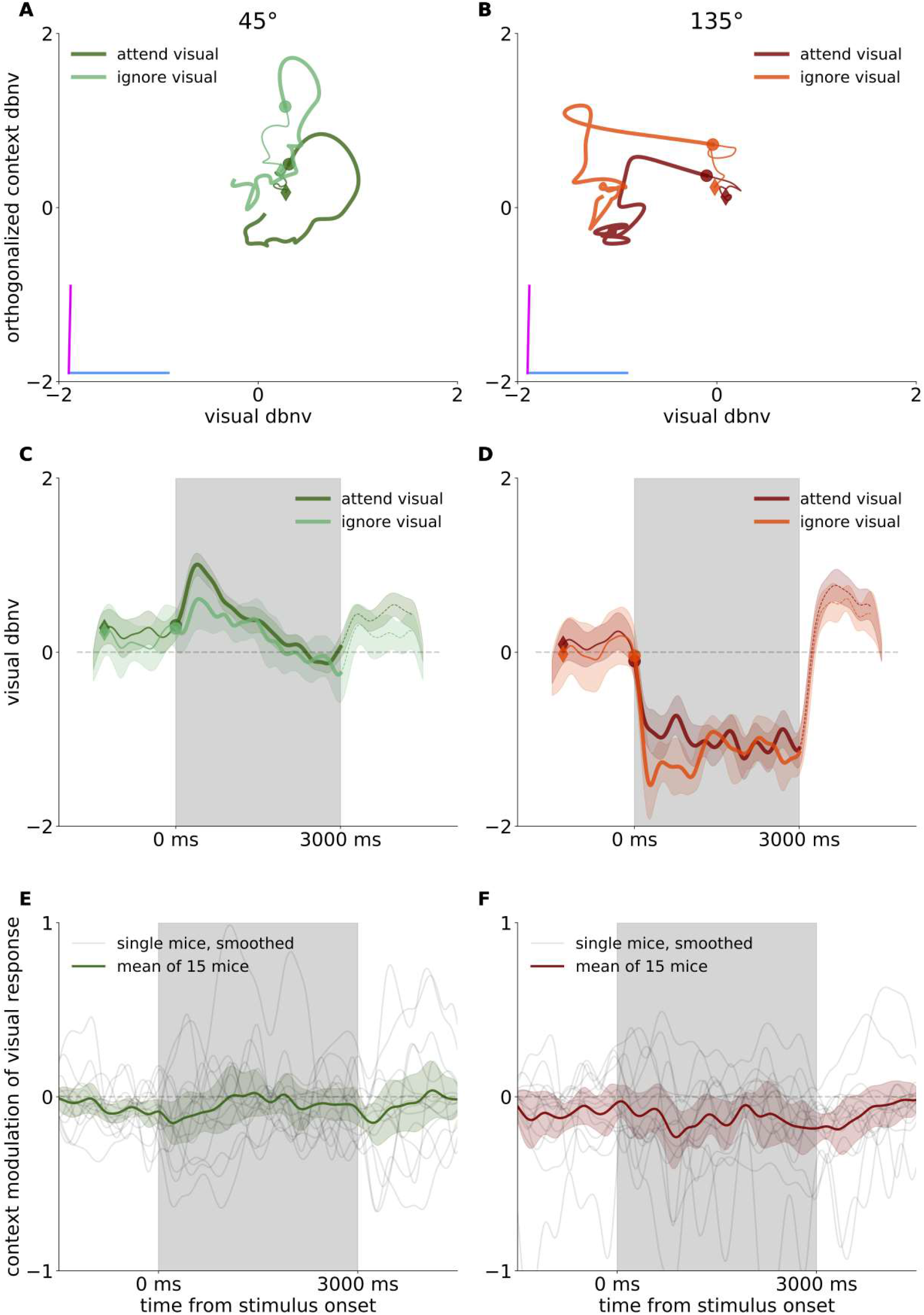
Effect of context on processing visual information. **A**, Time course of population trajectory projected on the subspace stretched by the context and visual DVs for trials with 45° grating stimuli. Data are only from trials with correct responses. *Dark* and *light colors* denote population trajectories in the attend visual and ignore visual contexts, respectively. Thin line shows activity in the pre-stimulus period while thick line shows activity in the on-stimulus period. Diamonds mark the start of the trajectory, while disks mark stimulus onset. **B**, Same as panel **A** but for 135° grating stimuli. **C**, Time course of the population response to the 45° stimulus projected on the visual DV. Colors show different task contexts, only correct trials are analyzed. **D**, Same as panel **C**, but for the 135° grating stimulus. **E-F**, Differences of visual DBNV-projected populations trajectories between different task contexts.

In order to analyze the activity dynamics directly related to the stimulus we projected the population activity on the visual DV (Fig. 5C, D). The time course of the population trajectory showed high stimulus content specificity but was close to identical in the two task contexts, indicating that the different initial conditions have marginal effect on the activity pattern related to the visual stimulus. We characterized the context-specificity of the visually elicited population responses by the difference of the context-conditioned responses. Repeating the analysis with the whole population of recorded animals, no context-specificity in stimulus-elicited responses could be established (Fig. 5E, F, dependent t-test p=0.48+/-0.01 and 0.19+/-0.01 for 45 and 135 stimuli respectively over the time course from stimulus start). In summary, despite distinct initial conditions in different task contexts, the dynamics of evoked population activity followed similar trajectories.

### Choice- and context-related activity in V1 are orthogonal

Beyond visual content and task context, both individual neurons and population responses showed sensitivity to the choice animals made. Choice-related activity was not expected to be independent from visual content related activity since the choice depended on the identity of the presented visual stimulus in the attend-visual context. In addition, choice-related activity was also difficult to differentiate from movement related activity since movement preparatory activity and even movement related activity could appear during the stimulus presentation period. Therefore choice related activity captured by decoding analysis did not solely convey information on the potential cognitive variable signalling the decision of the animal at later stages of a trial. Still, its relationship to task context provided an important insight into how the latent variable context is represented in relation to other task-relevant activity. Therefore, we constructed a linear decoder for choice (Fig. 6A). Choice could be decoded throughout the trial and also after stimulus offset. As expected, across-animal comparisons showed higher decodability later during the trial (Fig. 6B, accuracy early 0.63+/-0.02, late 0.70+/-0,02, one-sided dependent t test p=0.002, 15 animals). In order to establish the relationship of context and choice representations, we analyzed population responses in the two-dimensional subspace of the context and choice DVs (Fig. 6C). Population responses in this DV-subspace were clearly distinguished according to the choice the animal made and change in task context resulted in a close to orthogonal shift. Comparison of DV angles across all the animals in the recorded population confirmed a strong peak around ninety degrees, indicating that the task context is represented in a subspace orthogonal to choice-related activity (choice DVs assessed separately in attend: 86.2+/-2.5°, and ignore: 91.5+/-2.7°). Multiple sources contribute to choice-related activity, which includes both cognitive factors, such as reward expectation, and movement-related activity, such as preparation of action, locomotion, etc. These factors cannot be distinguished by our analysis but the orthogonality to the context DV demonstrates that choice-related activity comprises an independent source of variance in V1.

**Figure 6.**
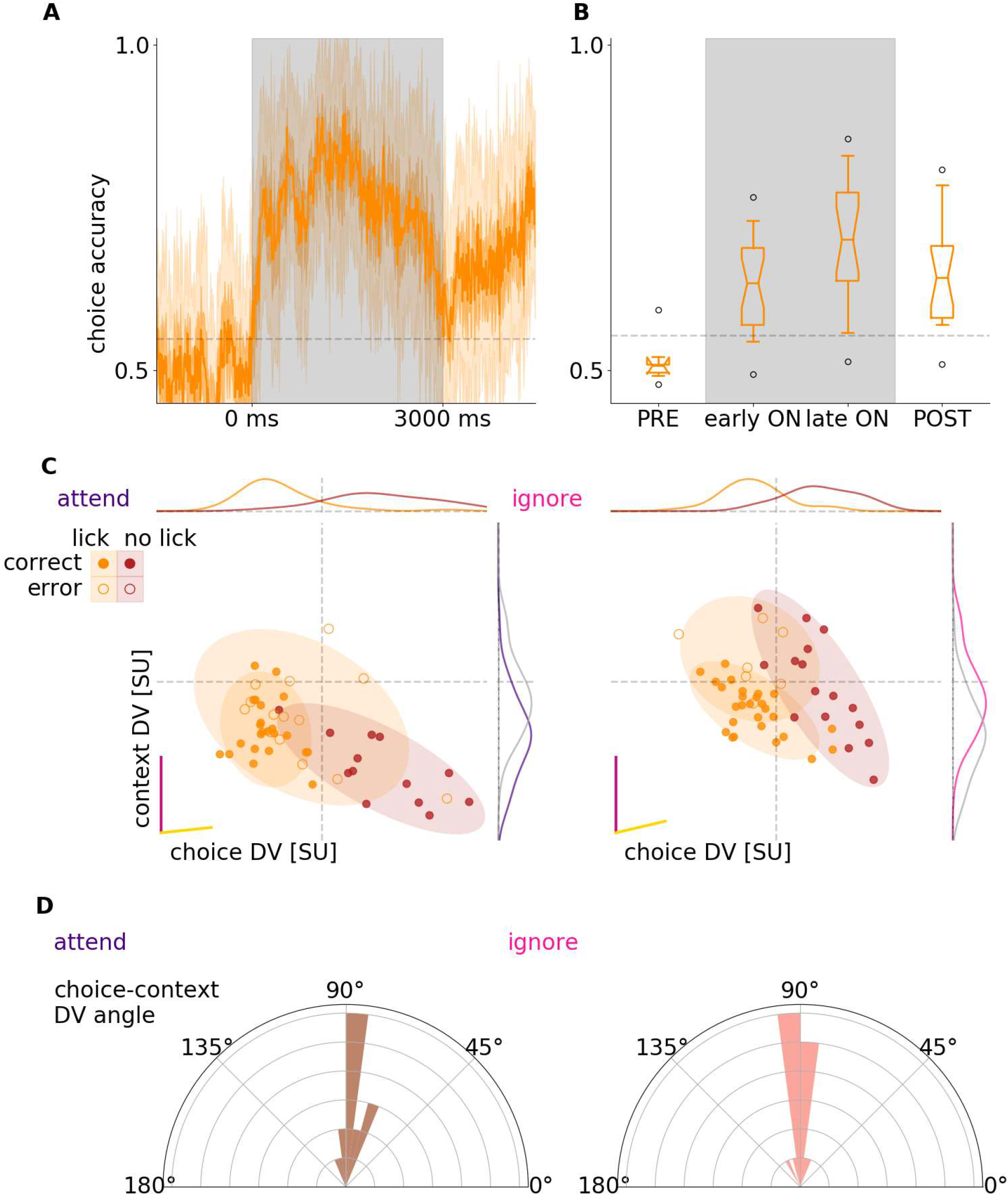
Choice-related activity. **A**, Decoder performance in 50 ms time windows at 10 ms sliding resolution for the choice the animal made in a trial (lick/no-lick), prior to stimulus onset (PRE), during stimulus (ON), and after stimulus (POST) for an example animal. Gray shading marks the stimulus presentation period. Grey dashed line indicates 1 s.e.m. of randomized labels decoder accuracies around chance level. **B**, Average performance of visual decoder for all animals. Box and whiskers denote 25-75, and 2.5-97.5 percentiles respectively, midlines are the means, notches are 95% confidence level error of the mean. Grey lines as in **B**, but averaged over animals. **C**, Population responses projected on the DV subspace in individual trials (dots) in different task contexts (*left* and *right panels*) and with different choices the animal made (*orange* and *red* for lick and no lick, respectively) for an example animal. Since choice decoder changes across task contexts, trials in different contexts are projected on their respective DV subspace (*left* and *right* panels for attend visual and ignore visual context, respectively). The two panels share the direction of the context DV (vertical axis). Magenta and yellow lines denote the DV directions of context and choice decoders, respectively. Horizontal distributions show the marginal of the responses on the choice DV. Vertical distributions show the marginal of population responses on the context DV with purple and pink lines corresponding to the marginals on the context on the panel and grey distributions showing marginals from the opposite panel. **D**, Histogram of the angle between context and choice DVs across animals in the two task contexts (*left* and *right*). **E**, Effect of choice on visual DV-projected population activity in an example animal **F**, Choice-related shift in visual DV-projected population activity for all animals.

### Decreased discriminability of visual information in error trials

In order to test whether choice had an effect on visual stimulus discrimination, we trained visual identity decoders on trials where only the choice differed but were identical otherwise. Separate decoders were trained from neural activity on subsets of go and no-go trials: Data was conditioned on the attend visual context only, and pooling all trials from all mice we separately assessed the accuracy for combinations of go and no-go signals, and correct and error trials. We have found no statistical evidence of different visual discriminability between hit and miss trials for the go signal (DV projected activity, t test p=0.059). Note that training to saturating performance yielded very few miss trials, which quenched statistical power. However, for no-go trials animal performance significantly altered discrimination decoding (t test p<10^−6^). In false alarm trials activity projected onto the visual DV were on average closer to the boundary than in correct rejection trials when trials from all mice were aggregated (Supplementary Fig 5A). Of the 15 mice, 7 individiual mice showed significantly worse discrimination in error trials, 3 mice significantly better, while in 5 mice, activity was not distinguishable based on behaviour (Supplementary Fig 5B). Modulation of the discriminability of the responses by behavioral outcome is an indication that a more reliable code might correspond to better behavioral performance.

### Controlling for movement related activity

In our paradigm, task context was stable in the first half of the session and was switched in the second half of the session. As a consequence, changes in movement patterns that occur between these periods can introduce confounds in our analysis of context representation since differences in movement-related activity could be picked up by the task context decoder.

To determine if the context signals we identified in population activity are true cognitive variables and not merely a result of changes in locomotion patterns, we devised a locomotion matching analysis. We used locomotion data to match the distribution of movements across task conditions similar to firing rate matching in Bányai et al. (2019) and mean matching in Churchland et al. (2010) (Materials and Methods). Briefly, we constructed a joint distribution of running speed measurements and population activity measurements in 50-ms time windows. Thus, any given time window in a trial yields a point in the multi-dimensional space of running speed and population activity. To reduce sampling noise in this distribution, we collected data from multiple consecutive time windows and data points from these time windows were collectively used to construct a single distribution. The resulting marginal run speed distribution is thus the union of noisy histograms collected from a longer period of time but individual points represent 50-ms time windows (Fig. 7A). Separate distributions were constructed for the two task conditions. In order to control for running speed differences in the two task contexts we randomly subsampled the points of the joint distribution such that the running speed histograms were matched across conditions (Fig. 7B,C). We compared context decoder accuracy differences between running speed matched and non-matched trials (Fig. 7D). We found no difference between running speed matched and non-matched decoders: differences are one order of magnitude smaller than the 2 s.e.m. range of cross-validations in all animals involved in the locomotion matching analysis (Fig 7E, average of means over the time course of accuracy differences from 6 animals 0.003+/-0.003, and standard deviation 0.021+/-0.003). To control for potential delayed effect of locomotion on V1 activity, we also performed the analysis with time shifts between the locomotion and neural response data. In addition to motion matching with no shift, 100 and 200-ms shifts were introduced both forward and backward in time. We found no effect of this shift on locomotion-matched context decoding performance (Fig 7F).

**Figure 7.**
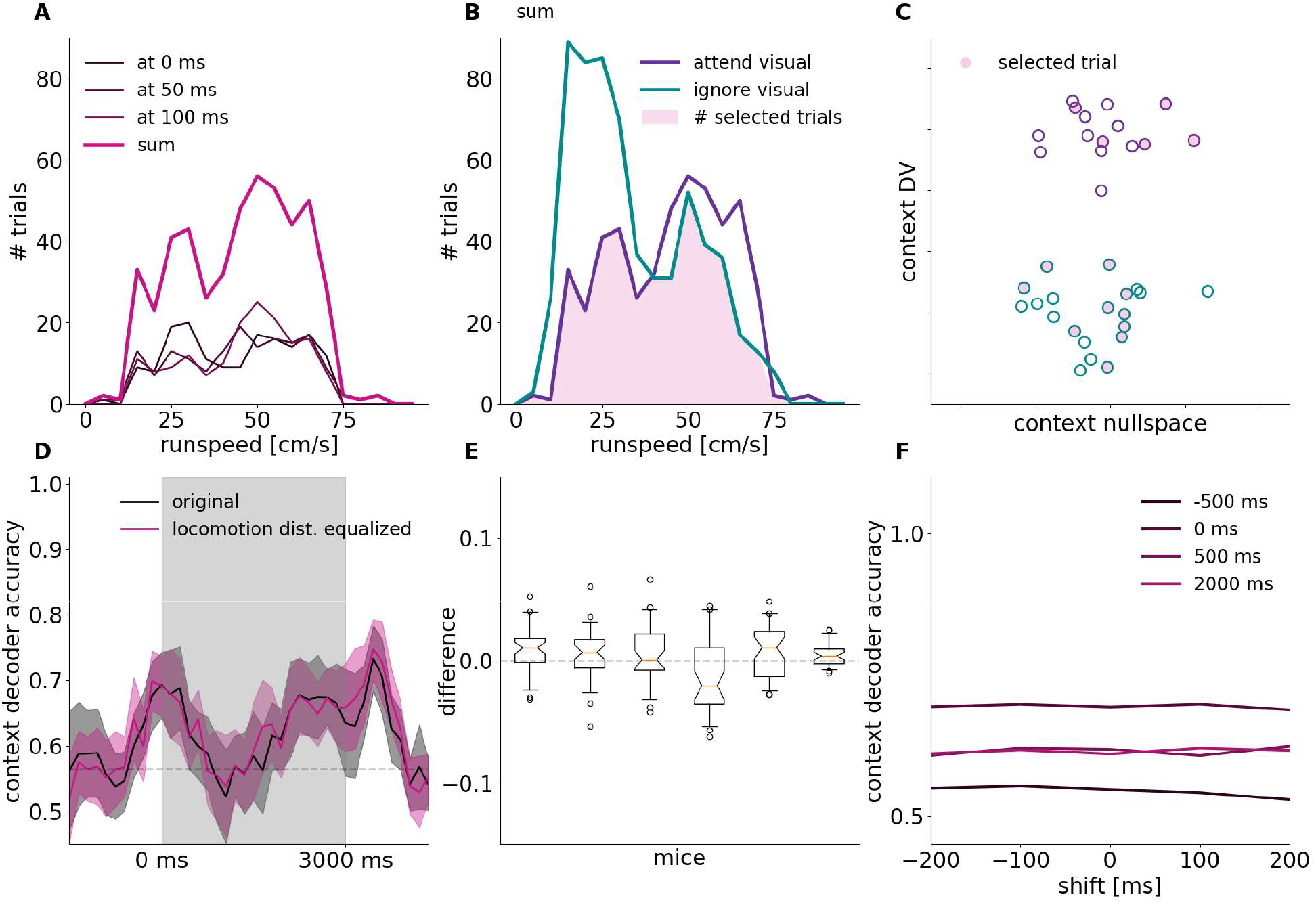
Controlling for differences in locomotion intensity between task contexts. **A**, Example histograms of running speed at different time points during the trial (*thin lines*) and summary histogram obtained by collating the three time-specific histograms (*thick line*). Resolution of runspeed histograms is 5 cm/s. Histogram was constructed for trials recorded in the attend visual task context. **B**, Task-context specific running speed histograms for the attend visual and ignore visual conditions (*purple* and *green lines*, respectively). Histograms are the summary histograms shown of panel **A**. Running speed distributions over trials in the attend visual and ignore visual conditions are matched by subsampling equal number of trials at each run speed bin (*shading*). **C**, Schematics illustrating run speed matching in the population activity space. Trial-by-trial population responses (*circles*) shown on a two-dimensional subspace, the vertical axis is aligned to the context DV. Subsampling trials based on the run speed distributions (panel **B**) yields a selection of trials in which the distribution of running speed is equal in the two task conditions (*filled circles*). **D**, Context decoder accuracy averages of 10 locomotion matched subsampling (magenta) and randomized control with same trial numbers (black). Filled area 2 s.e.m. of 10 fold cross-validation. Grey dashed line indicates 1 s.e.m. of randomized labels decoder accuracies around chance level. **E**, Difference between run speed-matched and control decoder accuracies from **D** in a population of mice, whisker plots indicate distribution over time course for each mouse. **F**, matched accuracies, same as **D**, at single points along the trial (colored lines), with run speed shifted relative to neural activity (horizontal axis).

Since distribution matching resulted in identical running speed contributions to neural activity in both contexts, differences in locomotion could not contribute to decoding the context representation from neural activity. This analysis eliminates the effect of potential V1 activity shifts caused by overall running speed differences (Dadarlat & Stryker, 2017). We can not rule out that other movement-related activity that are not detectable by measuring running speed could contribute.

## Discussion

We showed the existence of a neuronal population representation of a cognitive variable, task context, in V1. Importantly, task context was a latent variable since it was not directly cued by sensory stimuli and was therefore inferred by the animal through the contingencies between multimodal stimuli and water rewards. Representation of task context was ‘mixed’ with that of visual stimuli since overlapping populations showed sensitivity to both. Multi-dimensional subspace analysis revealed that the representation of task context was orthogonal to the subspace where visual grating stimuli were represented. Furthermore, despite engaging overlapping populations, task context did not affect the population dynamics induced by visual stimulus presentation and was therefore represented independently from visual stimuli. We found a strong signal associated with task context in the inter-trial interval as well (off-stimulus period) in addition to visual stimulation (on-stimulus) periods. While the strength of task context representation during on-stimulus and off-stimulus activities was strongly correlated, the representation underwent a transformation: stimulus onset was characterized by a strong dynamical shift in the linear subspace where context-related variability could be identified. Therefore, population activity in V1 not only reflects the visual stimuli presented to the animal, but also directly represents cognitive variables such as task context.

Flexible use of available information is critical for intelligent behavior (Niv, 2019). Depending on the context, the same sensory stimulus can invoke vastly different behavioral patterns. Based on behavioral output alone, previous studies have proposed that task context is represented through latent variables (Courville, Daw & Touretzky, 2006; Courville et al., 2004; Gershman & Niv, 2010). According to these accounts, learning about the dependencies between conditioned and unconditioned stimuli is achieved by introducing latent variables which correspond to the ‘rules’ of the task. These accounts are in contrast with traditional approaches of learning where ‘associations’ are learned between the conditioned and unconditioned stimuli, without the introduction of additional latent variables. The details of the neural representation of such local associations versus latent variables have remained elusive. Lesion experiments indicate that in rodents the orbitoforntal cortex can be critical for maintaining latent task variables such as context (Wilson et al., 2014). This idea gained further support from a human imaging study where orbitofrontal was shown to be central to representing the latent variables of the task (Schuck et al., 2016) but this study also showed some selectivity for certain latent task variables in the visual cortex. In light of these studies, our results suggest that the effects of the latent task context can be identified as early as the primary visual cortex. Such latent variable representations are necessary for efficient reinforcement learning (Niv, 2019) and are also critical components in forms of unsupervised learning, such as structure learning (Gershman, Norman & Niv, 2015; Orbán & Wolpert, 2011). Our findings provide insights into the properties of the representation of latent variables at the level of the primary visual cortex, yet the way the primary visual cortex contributes to task-dependent processing of sensory signals remains to be established.

Rule or context selective neural firing has been repeatedly observed in frontal and parietal cortices (Mansouri, Matsumoto & Tanaka, 2006; Stoet & Snyder, 2004; Wallis, Anderson & Miller, 2001; Wallis & Miller, 2003). Whether these neural dynamics impact firing in primary sensory cortices during context-dependent decision making is poorly understood. A study of prefrontal cortex (PFC) population dynamics during context-dependent decision making in non-human primates shows that the PFC receives unfiltered visual information which undergoes differential dynamics based on the appropriate context (Mante et al., 2013). The study argued against the alternate scenario where top-down input acts within V1 to select the contextually appropriate input and relays only those to PFC. While the design of our study is distinct in that it requires contextual information to be maintained across multiple trials, our results provide additional insight into this question: while contextual information shapes the activity in V1, it does so in a subspace orthogonal to the representation of relevant stimuli but leaves V1 dynamics in the subspace relevant to stimulus representation intact. Our work suggests that if contextual information represented in higher order areas is relayed back to V1, it is represented in an orthogonal population space. This is also consistent with previous work showing context-dependent changes in the stimulus selectivity of population responses in mouse V1 (Khan & Hofer, 2018; Poort et al., 2015) or context-dependent modulation of V1 firing during navigation (Saleem et al., 2018).

The context-dependent changes we identified in V1 population responses can be interpreted as a source of noise correlation since changes in context introduce correlated changes in population activity. Our results show that these changes are orthogonal to the stimulus dimension therefore do not contribute to information limiting correlations (Moreno-Bote et al., 2014). A recent study in non-human primates has highlighted that a feedback-driven component of noise correlation that is measured within a given task context changes across tasks and displays a structure reminiscent of information limiting correlations (Bondy, Haefner & Cumming, 2018). Thus, noise correlation measured within a task context and across task contexts might indicate different feedback components and their relationship can reveal the exact mechanism of context-dependent modulations (Haefner, Berkes & Fiser, 2016; Haimerl, Savin & Simoncelli, 2019). Such context-dependent changes in noise correlations are not constrained to V1 but can be identified in MT and IT as well (Cohen & Newsome, 2008; Koida & Komatsu, 2007; Tajima et al., 2017). Recently, two detailed analyses of the choice-related activity in the MT of non-human primates demonstrated that much of the choice related activity is orthogonal to the stimulus-induced variance rather than being aligned to it (Ruff & Cohen, 2019; Zhao et al., 2020). Our analysis is consistent with this view and along with the orthogonal representation of the context, highlights that task-relevant variables partition the activity space into orthogonal subspaces. In mice, a small population of PFC neurons have rule selective firing during the delay period of a cued context-dependent decision making task similar to the task in this paper, suggesting that the mouse PFC can encode context (Rikhye, Gilra & Halassa, 2018). Our task which requires the maintenance of contextual information shows that this information can be decoded from V1 neurons in a dynamic fashion across multiple trials.

We found that the context signal undergoes a transformation during the transition from off-stimulus to on-stimulus condition but remains relatively stable within the conditions. This observation can be puzzling since the contextual variable itself is invariant across on and off stimulus conditions on a larger timescale. In terms of the dynamics of neural populations, such transformation can be interpreted as changing fixed points that the network converges to in the two conditions. Indeed, recent analysis of recurrent dynamics in networks has started to uncover the relationship between learning principles, training schedule, and aspects of neural dynamics, such as condition-dependent fixed point dynamics, and low dimensional dynamics in independent subspaces (Duncker et al., 2020). In our study, the transformation of the context representation could not be traced back to a simple dynamics in which a subpopulation of neurons switch their activities between conditions since a set of neurons displayed stable contribution to the context decoder throughout the trial. Our findings show that while it is possible to read out from the population activity the task being performed throughout the trial with simple linear decoders but only a select number of neurons is appropriate to construct a decoder that works equally well during and before task execution. We show that for a more efficient read-out of task identity from the whole population a pair of decoders is more efficient.

How does contextual information reach V1? As V1 receives top-down inputs from anterior cingulate (Zhang et al., 2016), motor, premotor, retrosplenial (Makino & Komiyama, 2015), posterior parietal (Lyamzin & Benucci, 2018), and higher visual cortices (Pak et al., 2020), as well as ascending inputs from lateral posterior thalamic nucleus (Roth et al., 2016), there may be multiple direct and indirect inputs that could be converging upon V1 to give rise to contextual coding. Previous work has highlighted an indirect route from the medial prefrontal cortex to the basal ganglia onto perigeniculate reticular nucleus neurons which in turn inhibit lateral geniculate neurons projecting onto V1 (Nakajima & Halassa, 2017). This circuit can inhibit irrelevant visual information when the animal is basing its decisions on simultaneously presented auditory input, similar to the task in this paper (Wimmer et al., 2015). Other work has highlighted how anterior cingulate firing, typically on conflicting decisions (Botvinick et al., 1999), can either directly or indirectly through the superior colliculus modulate thalamic lateral posterior nucleus neurons which in turn modulate visual cortical neurons (Hu et al., 2019). Direct inputs from the auditory cortex may also modulate V1 firing (Deneux et al., 2019; Ibrahim et al., 2016; Meijer et al., 2017, McClure and Polack, 2019). Feedback from the multisensory-integrator posterior parietal cortex (PPC), which is below PFC in the hierarchy, may also influence V1 during task engagement as PPC neurons appear to respond only to task relevant stimuli (Pho et al., 2018). It is also possible that neuromodulatory inputs could play a role, either directly or through modulation of interneurons (Hattori et al., 2017).

V1 activity is strongly modulated by locomotion (Niell & Stryker, 2010; Polack, Friedman & Golshani, 2013). In fact, recent studies suggest that the majority of stimulus-unrelated activity in V1 is associated with locomotion and movement (Stringer et al., 2019a). Importantly, this study also implied that this stimulus-unrelated component of activity occupies the same activity subspace which is occupied by off-stimulus activity. This subspace shared a single linear dimension with stimulus-evoked activity but was orthogonal apart from this single dimension. Our findings demonstrate that beyond these movement-related signals, signatures of latent task context can also be identified in the off-stimulus activity. We found a similar orthogonal relationship between the subspace occupied by task context and that of visual stimuli (albeit we only used simple grating stimuli instead of more structured ones). In order to exclude the possibility that the task context signal identified in the analysis is contaminated by variations in locomotion patterns, we performed control analysis where locomotion statistics was matched across task context, and confirmed that the identified task context representation was unaffected by variations in locomotion patterns. Previous studies have also shown that a wide range of uninstructed movements occur during task execution (Musall et al., 2019) and variations in fine-scale movements contribute to activity patterns in the cortex, including V1. Our control for motion excludes larger scale features of uninstructed movements contributing to the context representation.

It will be important to identify the inputs to V1 that generate the contextual representation. This will require recordings from neurons projecting to V1 during the task as well as manipulations of their activity patterns during off-stimulus and on-stimulus periods. Chronic recordings from large populations of cortical neurons during both learning and performance of the task will be critical for understanding how these representations emerge and are maintained across days. Our results show that V1 activity is systematically modulated by the task the animal is engaged in. Recent behavioral explorations have demonstrated that the rule that the animal bases its decisions on can change during a behavioral session, even when the task structure remains unchanged (Roy et al., 2020). It will be important to see whether such spontaneous changes in behavioral strategy are reflected in V1 activity in a similar manner to how changes in task context are represented. Finally, it will be important to identify how the contextual code in V1 contributes to perception and decision making in conditions at the edge of the perceptual threshold.

## Acknowledgements

The authors thank Rachna Goli, Allison Foreman, Brandon Sedaghat, and Tara Shooshani for their help with training the animals to perform the behavioral task. The authors would also like to thank Anne Churchland and Dario Ringach for their comments on a previous version of the manuscript. P.G. and G.O. were supported by a grant from the Human Frontiers Science Program, P.G. was supported by grants 1R01MH105427, R01NS099137, 1P50HD103557, M.A.H. and G.O. were supported by a grant by the Hungarian Brain Research Program (2017-1.2.1-NKP-2017-00002).

## Author Contributions

P.O.P., M.E, and P.G. designed and optimized the behavior, and designed all experiments. D.T and M.E. trained animals and performed the recordings. D.T, M.V.M. and K.S. curated the data. M.A.H. and G.O. designed the analysis. M.A.H. performed the analysis mainly with input from G.O., but also P.O.P. and P.G. M.A.H., P.O.P, P.G. and G.O. wrote the manuscript.

## STAR Methods

### SURGERY

All experimental procedures were approved by the University of California, Los Angeles Office for Animal Research Oversight and by the Chancellor’s Animal Research Committees. 7-10 weeks old male and female C57Bl6/J mice were anesthetized with isoflurane (3–5% induction, 1.5% maintenance) ten minutes after intraperitoneal injection of a systemic analgesic (carprofen, 5 mg/kg of body weight) and placed in a stereotaxic frame. Mice were kept at 37°C at all times using a feedback-controlled heating pad (Harvard Apparatus). Pressure points and incision sites were injected with lidocaine (2%), and eyes were protected from desiccation using artificial tear ointment. The surgical site was sterilized with iodine and ethanol. The scalp was incised and removed, and a custom-made lightweight omega-shaped stainless steel head holder was implanted on the skull using Vetbond (3M) and dental cement (Ortho-Jet, Lang), and a recording chamber was built using dental cement. Mice recovered from surgery and were administered carprofen for 2 days, and were administered amoxicillin (0.25 mg/ml in drinking water) for 7 days. Mice were then water-deprived and trained to perform the behavior (discussed below).

Approximately 24 hours before the recording, mice were anesthetized with isoflurane, a small craniotomy (0.5 mm diameter) was made above the right cerebellum and a silver chloride ground wire was implanted within the craniotomy and fixed in place with dental cement. A circular craniotomy (diameter = 1 mm) was performed above the right V1 (V1 targets were determined by regression of adult brain lambda-bregma distances: 1.7–2.5 mm lateral and 0.0 – 0.5 mm rostral to lambda. The exposed skull and brain were covered and sealed with a silicone elastomer sealant (Kwik-Sil, WPI). On the day of the recording, the mouse was placed on the spherical treadmill and head-bar fixed to a post. The elastomer sealant was removed and the craniotomy chamber was filled with cortex buffer containing 135 mM NaCl, 5 mM KCl, 5 mM HEPES, 1.8 mM CaCl_2_ and 1 mM MgCl_2_.

### BEHAVIORAL TRAINING

Following implantation of the headbars, animals recovered over 3 days, and received 10 to 20 minutes of handling per day, thus habituating the animals to human interaction for 4 days. Animals were then water-deprived, receiving approximately 1 mL of water per day. During this time, animals were placed on an 8-inch spherical treadmill (Graham Sweet) in the behavioral rig for at least 3 days to habituate to head-fixation for 15 minutes per day. The spherical treadmill was a Styrofoam ball floating on a small cushion of air allowing for full 2D movement (Graham Sweet, England). The animal’s weight was measured daily to ensure no more than approximately 10% weight loss.

Animals were first trained to perform unimodal visual and auditory lick/no-lick (go/no-go) discrimination tasks. Licks are detected by using a lickometer (Coulbourn Instruments). Lick detection, reward delivery and removal, sensory stimulation and logging of stimuli and responses were all coordinated using a custom-built behavioral apparatus driven by National Instruments data acquisition devices (NI MX-6431) controlled by custom-written Matlab code. A 40-cm (diagonal screen size) LCD monitor was placed in the visual field of the mouse at a distance of 30 cm, contralateral to the craniotomy. Visual stimuli were generated and controlled using the Psychophysics Toolbox (Brainard, 1997) in Matlab. In the visual discrimination task, drifting sine wave gratings (spatial frequency: 0.04 cycles per degree; drift speed: 2Hz; contrast: 100%) at 45 degrees, moving upwards, were paired with a water reward. Drifting gratings of the same spatial frequency but at 135 degrees orientation, moving upwards, signaled a reward would not be present, and the animal was trained to withhold licking in response to the stimulus. The inter-trial interval was 3 seconds, except for trials in which the animal had a miss or false alarm, then the inter-trial interval was increased to 6.5 seconds. The animal’s behavioral performance was scored as a d’ measure, defined as the z-score of the hit rate minus the z-score of the false alarm rate. Once animals reached expert performance (d’>1.7, p<0.001 as compared to chance performance, Monte-Carlo simulation), they were advanced to learning the auditory discrimination task where a low pure tone (5 kHz, 90 dB) indicated that the animal should lick for reward and a high tone (10 kHz, 90 dB) indicated that the animal should withhold licking. The inter-trial interval was similarly 3 seconds and the inter-trial interval was increased to 9 seconds after misses or false alarms. After animals learned the auditory discrimination task (d’>1.7) they were trained to perform the multimodal attention task. In this phase, animals first performed one block of visual discrimination (30 trials). If their performance was adequate (d’>2.0, correct rejection rate>70%, hit rate>95%) they then performed the visual discrimination task with auditory distractors present (the high or low tones) for 120 trials. Then, after a five-minute break, they performed the auditory discrimination task for 30 trials and if their performance was adequate (d’>2.0, correct rejection rate>70%, hit rate>95%), they performed auditory discrimination with visual distractors present (oriented drifting gratings at 45 or 135 degrees described previously). During each training day and during the electrophysiological recordings, each trial set started with 30 trials where only visual or auditory stimuli were delivered which signaled whether the animal should base its decisions on the later multimodal trials to visual or auditory stimuli respectively. Each trial lasted 3 seconds. When the cue stimulus instructed the animal to lick, water (2 microliters) was dispensed two seconds after stimulus onset. No water was dispensed in the no-lick condition. To determine whether the animal responded by licking or not licking, licking was only assessed in the final second of the trial (the response period). If the animal missed a reward, the reward was removed by vacuum at the end of the trial. Animals performed 300-450 trials daily. Only one training session was conducted per day with the aim to give the animal all their daily water allotment during training. If animals did not receive their full allotment of water for the day during training, animals were given supplemental water one hour following training. Whether the animal started with attend-visual or ignore-visual trial set was randomized. Importantly, the monitor was placed in the exactly the same way during the auditory discrimination task as it was placed during the visual discrimination task, and a grey screen, which was identical to that during the inter-trial interval of the visual discrimination task and isoluminant to the drifting visual cues, was displayed throughout auditory discrimination trials. As a result, the luminance conditions were identical during visual and auditory discrimination trials.

### MOTION DETECTION

Head-fixed animals run on a treadmill consisting of an 8-inch Styrofoam ball (Graham Sweet) suspended 2mm above an 8.5-inch Styrofoam cup (Graham Sweet) using pressurized air. Mouse treadmill rotation was recorded as an analog signal, using a custom printed circuit board based on a high sensitivity gaming mouse sensor (Avago ADNS-9500) connected to a microcontroller (Atmel Atmega328). The signal was initially recorded along with electrophysiology at 25 kHz, then down-sampled to 1 kHz, and low-pass filtered < 1 Hz with a first order Butterworth filter. The processed signal was treated as a proxy of velocity. 6 animals were available for running speed measurements.

### IN-VIVO ELECTROPHYSIOLOGY RECORDINGS

Extracellular multielectrode arrays were manufactured using the same process described previously (Shobe et al., 2015). Each probe had 2 shanks with 64 electrode contacts (area of each contact 0.02 μm^2^) on each shank. Each shank was 1.05 mm long and 86 μm at its widest point and tapered to a tip. Contacts were distributed in a hexagonal array geometry with 25 μm vertical spacing and 16-20 μm horizontal spacing), spanning all layers of V1. Each shank was separated from the other 400 μm. The electrodes were connected to a headstage (Intan Technologies, RHD2000 128-channel Amplifier Board with two RHD2164 amplifier chips) and the headstage was connected to an Intan RHD2000 Evaluation Board, which sampled each signal at a rate of 25 kHz per channel. Signals were then digitally band-pass-filtered offline (100 – 3000 Hz) and a background signal subtraction was performed (Shobe et al., 2015). To ensure synchrony between physiological signals and behavioral epochs, signals relevant to the behavioral task (licking, water delivery, visual/auditory cue characteristics and timing, and locomotion) were recorded in tandem with electrophysiological signals by the same Intan RHD2000 Evaluation Board.

After performing the visual and auditory cross modal tasks, drifting gratings were presented to map the orientation selectivity of each recorded cell. A series of drifting gratings of 6 orientations spaced by 30 degrees, with both directions, randomly permuted, temporal frequency = 2 Hz, spatial frequency = 0.04 cycle per degree, contrast = 100%) was presented for 3 seconds, with a 3 second inter-trial interval, with each different presented 10 times.

### ACUTE MICROPROBE IMPLANTATION

On the day of the recording, the animal was first handled and then the headbar was attached to head-fix the animal on the spherical treadmill. The Kwik-Sil was removed and cortex buffer was immediately placed on top of the craniotomy in order to keep the exposed brain moist. The mouse skull was then stereotaxically aligned and the silicon microprobe coated with a fluorescent dye (DiI, Invitrogen), was stereotaxically lowered using a micromanipulator into the V1 (relative to lambda: 1.7–2.5 mm lateral and 0.0 – 0.5 mm rostral). This process was monitored using a surgical microscope (Zeiss STEMI 2000). Once inserted, the probe was allowed to settle among the brain tissue for 1 hr. Recordings of multiple single-unit firing activity were performed during task engagement (approximately 1 hr). After the recording, the animal was anaesthetized, sacrificed, and its brain was extracted for probe confirmation.

### DATA ANALYSES

#### Single unit activities (SUA)

Spike sorting was performed by Kilosort 2 (Pachitariu et al., 2016), and then manually curated in phy2 using MATLAB and PYTHON to yield single unit activities, with consistency criteria for autocorrelograms, interspike interval histograms, waveforms, maximal amplitude electrode locations, low false positive or missed spikes and also stable feature projections throughout the recording session. Highly similar clusters were merged manually if cross-correlation revealed identical refractory periods and if interspike interval histograms and feature distributions matched to provide a resulting unit without drift signs. Clusters were split when PCA feature space and interspike interval histogram showed mixtures of stationary distributions, and the cross-correlograms improved. Putative excitatory and inhibitory cell types were distinguished based on spike waveform characteristics corresponding to broad and narrow spiking. We applied a gaussian mixture model to cluster in the two dimensions of time from trough to peak and amplitude ratios of trough and peak on all cells from all mice.

#### Exclusion criteria to control for drift

We took great care such that only units showing no statistically identifiable drift in firing responses were included in the analysis. For this, signal to noise ratios of spike events were tracked during the parts of the recording session when the animal performed the task. For any given unit, drift was identified using a set of criteria on a long timescale. Specifically, we assessed the quality of the unit based on A) unit separability in feature space, B) stationarity of signal to noise ratio (S/N) and C) stationarity of firing rate. We excluded units that did not meet our criteria.

#### Spike counts

Spike counts were calculated in sliding windows in 10-ms bins and smoothed using a symmetric Gaussian kernel (σ=100 ms, optimized for linear decoding and typical firing rates). The kernel method approximates single trial instantaneous rate (IR). IRs were used in two distinct ways. First, we used IRs to assess absolute instantaneous firing rates (FR) in Hz. Second, for methods requiring standardized input, IRs were transformed to z-scores, with mean calculated from prestimulus (from −1500 ms to 0 ms) time-averaged baseline activities, while standard deviation was calculated for the whole trial. Sensitivity of single units to task variables was assessed by the largest mean rate difference between trial averages during different task epochs.

#### Neural activity vector

(referred to as population activity or activity) is defined as baseline-standardized SUA IRs as components of a time-varying vector. Neural activity vector space, specifically using the above coordinate system as basis, is defined as the space of possible activity patterns the neuron population can take (Fig 2A). A single point in this neural activity vector space corresponds to IR for each neuron, while trajectories describe dynamics of activity.

#### Relative signal variance

Total variance for signals with multiple conditions can be decomposed into the sum of noise variance and signal variance. Noise variance is the expected value over the conditions of the variances of the signal at specific conditions. Signal variance is the variance over the conditions of the expected values of the signals at specific conditions.

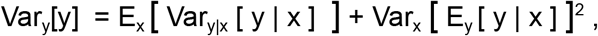

where x runs over two values of the condition, e.g. 45 and 135 degrees visual stimulus, and y runs over the trials with y | x signifying trials conditioned on the indexed values of x, e.g. only 45 degrees trials. Relative signal variance is a ratio of the signal variance and total variance, i.e. the second term on the right hand side divided by the term on the left hand side of the equation.

#### Principal Component Analysis

We concatenated neural activity of all time points along a trial for all trials, capturing both trial to trial and within trial variance. Principal Component Analysis was performed to obtain orthogonal directions by ordered variances of population activity. Neural activity was projected to various subspaces defined by a number of principal components (PC) by dot products with PC unit vectors. Relative variances were shown cumulatively for activities projected on a growing subset of PCs.

#### Linear decoding

We regressed neural activity with task variables using predictive trial-based cross-validation. Input to the decoder was a matrix with observations in different rows and features in columns. Observations were trials, each labeled with task variables such as stimulus identity, task context, and choice of the animal (licking or witholding licking). These identified nominal variables each consistently grouped trials into their respective two classes. Specific decoders were trained by conditioning on some of the non-decoded variables: for instance, on Fig. 5 we only selected successful trials for training the decoder. Feature space representing activity of all units in a time-segment was constructed by concatenating for each unit 5 consecutive 10 ms width spike count data points spanning 50 ms, and concatenating these vectors for each unit. The time resolution of the IFR kernel slide and the time-width of the decoder were optimized to saturate information transfer from raw data while preserving high time resolution. Classification of trials was trained with logistic regression. Separate decoders were trained for different time points of a trial with 10 ms resolution. Decoders used all single unit activities available, with the expectation that discrimination-irrelevant features would be averaged out with a Gaussian noise model in the log odds space. Unless otherwise noted, all decoders were performed with class-stratified 10-fold cross-validation. Decoder accuracy figures show means and two standard errors of means of the 9 CV test runs for each timepoint. Averages across mice only use the means, boxplots show animal population mean and 2 s.e.m. for notches and 25-75, 5-95 percentiles for the boxes and whiskers.

Cross-tested decoders were introduced to assess how a decoder trained in one condition performs in another condition. A decoder is trained at one particular time point of the trial and its performance is tested at a different time point. These time point crosstested decoders were trained with a cross validation scheme of 2 times 3:1 randomized train test splits. These were accurate enough, but consumed significantly less computation resources than full 10-fold CVs. Accuracy decay rate was calculated at each training point by fitting a linear function on the decoder performances in consecutive time points of cross-tested decoders and taking the slope over the first 500 ms of forward test shifts. Angles between time course-shifted (t1-t2) decoders were calculated as γ_t1,t2_ = arccos **d**^(t1)^**d**^(t2)^, **d** normalized at all t-s. The dimension of **d**-s when finding orthogonal vectors are irrelevant as we assume the nonimportant directions are random, thus highly likely to contribute zeros to the scalar product, as well as their noises cancel out to 0. Thus interesting angles are indistinguishable in N and 2 dimensional spaces. Average angles were calculated for each mice by averaging over the angle values in the lower triangles of prestimulus (−1500 to 0 ms) and on stimulus (0 to 3000 ms) block matrices, and the rectangular cross matrix (t1=-1500 to 0 ms, t2=0 to 3000 ms). To reduce overlapping effects within the instantaneous firing rate kernel width (100 ms) and the feature space width of 50 ms, angle-matrix elements closer to 100 ms from the diagonal (|t1-t2|<100 ms) were discarded.

We calculated effective chance levels for the data by averaging over 40 independent decoder crossvalidation accuracy distributions as described above, with fully randomized trial labels for the multimodal blocks taken from a Bernoulli(p=0.5) distribution. Thresholds indicate trial time course-averaged mean + 1 s.e.m of 40 runs of the means + 1 s.e.m.-s calculated over CVs. Decoders are expected to differ from chance only towards higher accuracy, thus the confidence level is one-sided 84% for 1 s.e.m. These were shown throughout multiple figures and panels as grey levels.

#### Assessment of the contributions of neurons to the decoder

The number of units recorded and successfully identified, and units that do contribute to the performance of a decoder varied across animals. Accuracy of decoders generally increases with the number of available neurons (Stringer et al., 2019b). We used partial correlation when comparing accuracies of separate decoders across animals to control for the number of units available. We performed linear least squares fit for the number of neurons predicting accuracy of decoders (Supplementary Fig. 2A,B). The residuals of these fits contain no information about the correlation between number of neurons and the accuracy of decoders. When such residuals of decoders that were fit to different conditions in the same animal correlated with each other (Supplementary Fig. 2C), the resulting coefficient is highly unlikely to be confounded by the number of neurons.

#### Decoder weight threshold between coding and noise ceiling

We used coefficient value-weighted histogram of coefficient values to determine which elements of the sum of products of predictor neuron activities and decoder coefficients are likely to contribute significantly to the decision vector direction on a trained decoder. We established a one order of magnitude threshold of signal to noise ratio, i.e. the lowest 10% of the weighted coefficients probability mass was regarded as noise. The decoder coefficient value corresponding to this 10 percentile we used as threshold, for 5 animals higher than 25 recorded neurons was 0.03. This threshold was robust to adding any and all other animals with lower number of units.

#### Multidecoder subspace analysis (MDSA)

A linear two-way classification decoder, e.g. logistic regression, defines the Decision Vector (DV) as the optimal one-dimensional subspace of the neural activity vector space along which neural activity realizations in single trials are best separated for the classes in question, e.g. 45 and 135 degree visual stimuli trials. In other words the DV is the vector perpendicular to the N-1 dimensional hyperplane that separates best the two classes (also called the normal vector of the decision boundary). We calculated DVs by averaging over the 10 cross-validation runs, and averaging the DVs of a number of linear decoders trained at different time points of the trial: Unless otherwise noted, DVs are time averages of the first 1500 ms of the trial after stimulus onset.

Since the coordinates of all decision vectors are in the same neural activity vector space, their angular difference, γ_1,2_ = arccos **d**^(1)^**d**^(2)^, where **d**^(k)^ is the normalized DV for the k-th decoder, provide a meaningful description of neural activity regarding task relevant variables the decoders were trained to discriminate.

Since decision vectors, **d**^(k)^ are defined in the same space (the population activity space), individual decision vectors can be combined together as a new basis spanning a multidimensional task-relevant subspace, *S*:

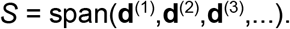

This subspace will contain task-relevant neural activity the linear decoders are able to pick up, and generally will be low-dimensional in our experimental paradigm. These bases do not necessarily form an orthogonal coordinate system, though: the basis vectors are linearly independent, but not necessarily orthogonal. Such skewed coordinate systems appear when some task variables have dependencies between each other, e.g. correct animal choice and stimulus identity of the relevant modality should be heavily dependent on each other if the animal performs the task well. We normalized the decision vectors to show each coordinate with unit length bases (SU, standardized units).

A low-dimensional representation of the high-dimensional neural population activity can be obtained by projecting the population activity onto the task relevant subspace. The relevant bases can include both a set of principal components (PCs) of a PCA, or subspaces of single or multiple DVs as basis:

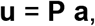

where **a** is the N dimensional activity vector, **u** is K dimensional subspace-specific activity, where K is defined by the number of PCs or DVs used as basis, and **P** is a K by N projection operator. **P** was defined by the K rows of transposed PC or DV vectors, respectively. Projection operators were normalized. We projected either each time point individually onto the projection bases, yielding full trajectories: **u**(t) = **P a**(t), or projected time-averages of the first 1500 ms of a trial: **u** = **P** <**a**(t)>_t_. Then the resulting task-relevant activity projection, **u**(t) or **u**, was conveniently visualized in an orthonormalized version of the coordinate system of the subspace *S* by the Gram-Schmidt procedure over **P** showing the first few basis in **P** as orthogonal as possible. The order of the bases was reordered for each question we asked to project the most relevant variables examined onto the most natural first few orthogonal coordinates.

When decoding from neural activity projected onto a subspace, the projection operator in the coordinates of the original neural activity basis is: **P** = **D** (**D**^T^**D**)^−1^ **D**^T^, where **D** = [ **d**^(1)^, **d**^(2)^, **d**^(3)^, …], a basis for *S*. In case of the 1 dimensional subspace of the visual decision vector, **D** = **d**, the projection **D** (**D**^T^**D**)^−1^ **D**^T^ will become **d d**^T^ **/** ‖**d**‖^2^.

#### Locomotion distribution matching

In order to separate the possible contribution of locomotion from the contribution of cognitive components to the context signal, we performed run speed distribution matching between contexts. We calculated the running speed of animals from the discrete time derivatives of the magnitude of the vectorial sum of forward and lateral displacement of the spherical treadmill. Running speed histograms were constructed between 0 and 100 cm/s at 5 cm/s bins. We averaged the run speed in 50-ms time windows, matching the length of the time window used by the linear decoders. The run speed data along with the population activity data defined a joint distribution in which every single trial in a context contributed a single data point. To control for differences between contexts, the histogram of run speeds in a given context served as an approximation of the distribution of run speeds. This histogram, however, is a noisy approximation since the set of approximately 60 data points that corresponds to the individual trials in a given context suffers from sampling noise. To rectify this issue we expanded the set by including data points from consecutive time windows. As a consequence, we could provide a more reliable estimate of speed distribution albeit in the end at a lower time resolution of 150 ms. To have a matching running profile in the two contexts, we subsampled the running speed distributions such that the histogram was identical in the two contexts. Then using the trials retained after subsampling, we performed the decoding for the context variable. We found that the increase in the number of trials using three consecutive 50 ms intervals balances out the number of trials we lose in all run speed bins, either from one or the other condition so that the decoders work on similar numbers of trials as without this matching on previous analyses. We averaged 10 of such matching subsamplings, and compared the cross-validated accuracy with the average accuracy of 10 randomized sub-samplings of the same number of trials for each run speed bin. Decoder results were such subsampling averages of means and 2 s.e.m.-s of the cross-validation accuracies. Shifts were defined as difference between time indices within trials between runspeed and neural data, unsyncing twe two, and the new full overlapping windows were used for analysis.

## Supplemental Information

**Supplementary Figure 1.**
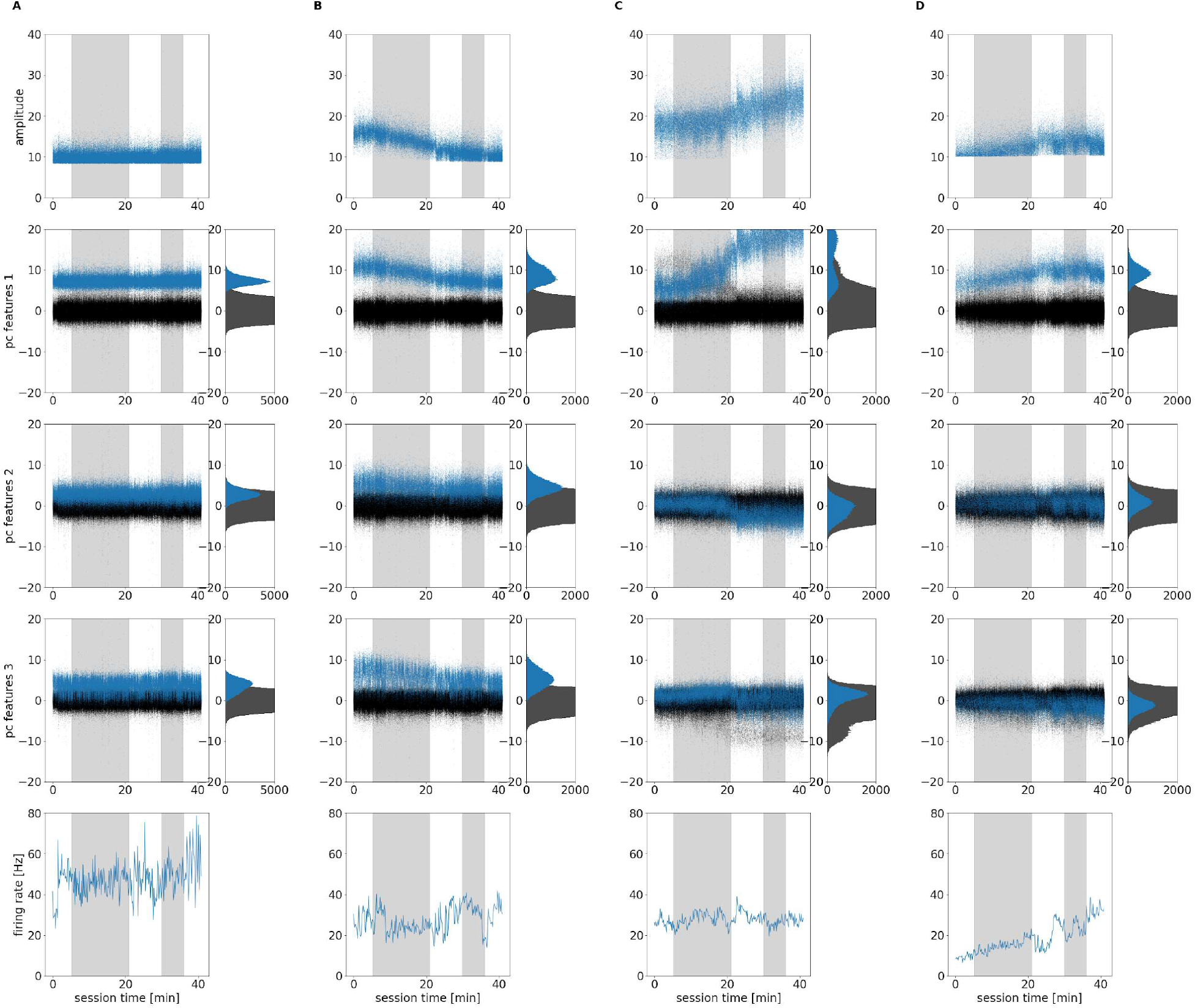
Selecting non-drifting units. Example spike characteristics as raw kilosort2 outputs along the recording session (horizontal axis) for trivial and edge case SUAs. *Columns* **A**-**D** are different SUAs. *Top row*: maximum amplitude of waveforms for each spike. *Rows 2,3,4*: wave projections onto the first three PCs of the best channel for the unit (blue dots) and all other units as background noise (black dots); *right side* on each panel: marginal histograms, counted from the total of the whole session in bins width of 0.4 standard principal component units. *Bottom row*: firing rate in 10 seconds bins. Grey shaded areas represent the multimodal stimuli blocks of the first and the second context. **A**, stable SUA, **B**, acceptable non-stationarity above noise ceiling with non-correlating firing rate, **C**, acceptable firing rate, but drifting in the first part of a cluster merge, **D**, visible drift. Drifting SUAs were discarded from further analysis.

**Supplementary Figure 2.**
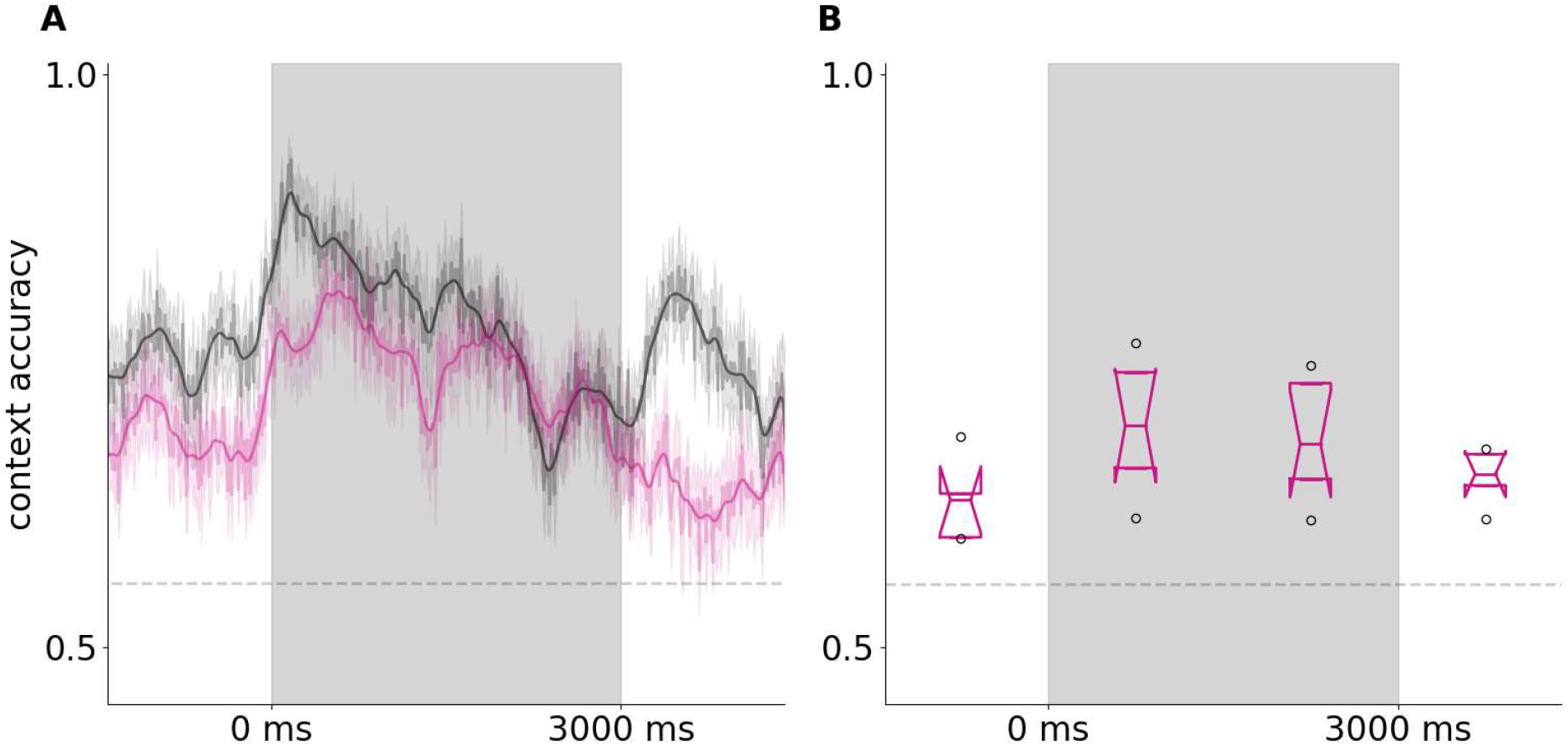
Context representation decoded from broad spiking neurons only. **A**, Context decoder accuracy timecourse using all available units, smoothed from Fig 3E (black line) and using only broad spiking units (magenta) on an example animal. **B**, Time-averaged mouse-population means of context representation using broad spiking units only, from a cohort of 5 animals, where all units were > 30 and broad spiking were >15. Grey lines are randomized chance levels as in Fig 3.

**Supplementary Figure 3.**
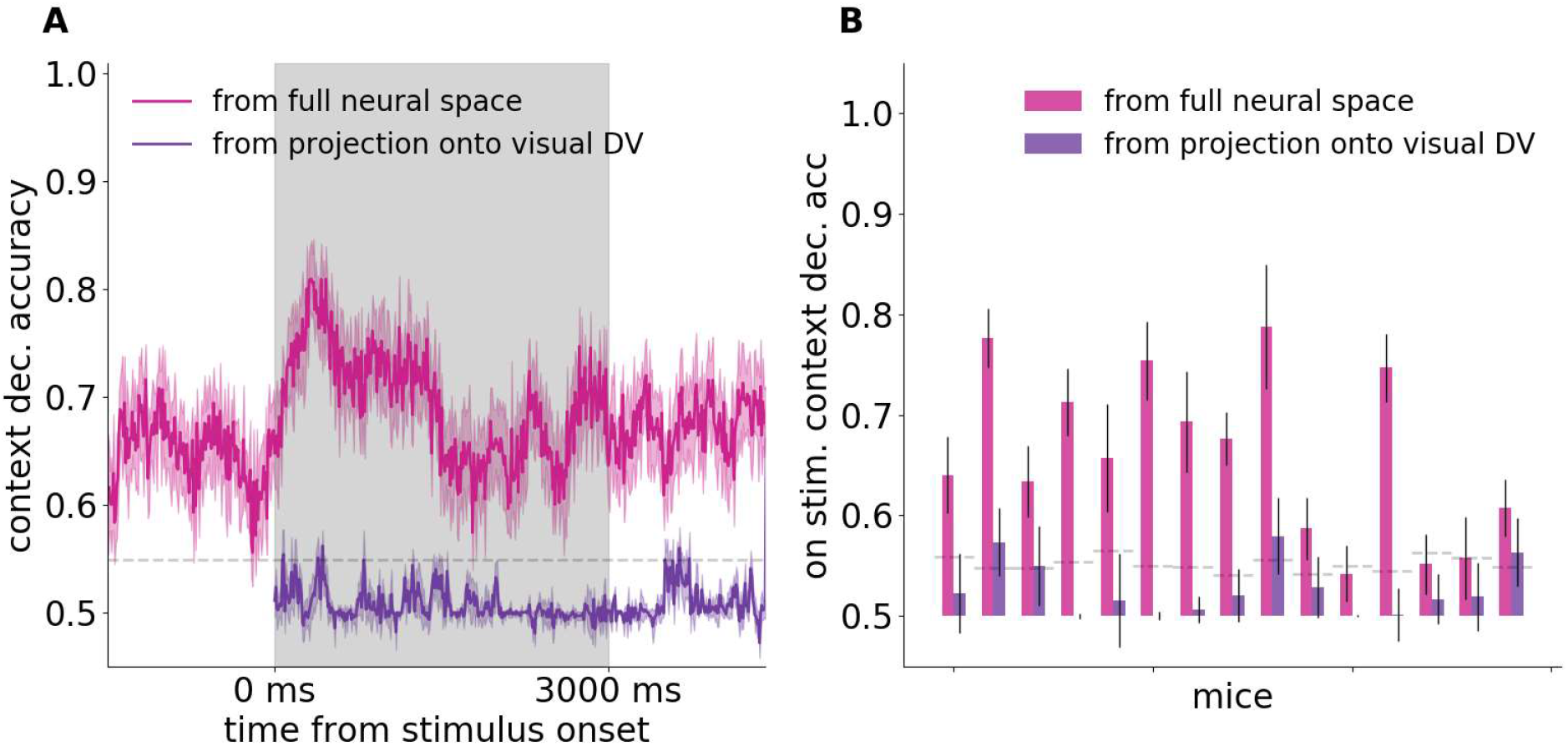
Context decoding from neural activity projected onto the visual decision vector subspace. **A**, Context accuracy timecourses of an example animal, decoding from the full neural activity space (magenta), and from neural activity projected onto the subspace defined by the visual decision vector (purple) at each timepoint along the trial. Solid lines are crossvalidation averages, with faint bands 2 s.e.m. Grey lines as in Supp Fig 2. **B**, For all mice separately (bar pairs, horizontal axis) time-averaged CV-mean accuracies during stimulus presentation, decoding from the full neural activity space (magenta bars), and from neural activity projected onto the subspace defined by the visual decision vector (purple bars). Error bars represent 1 standard deviation of the distribution of accuracies along the 300 timepoints. Grey lines representing randomized chance level are independently calculated for each mice.

**Supplementary Figure 4.**
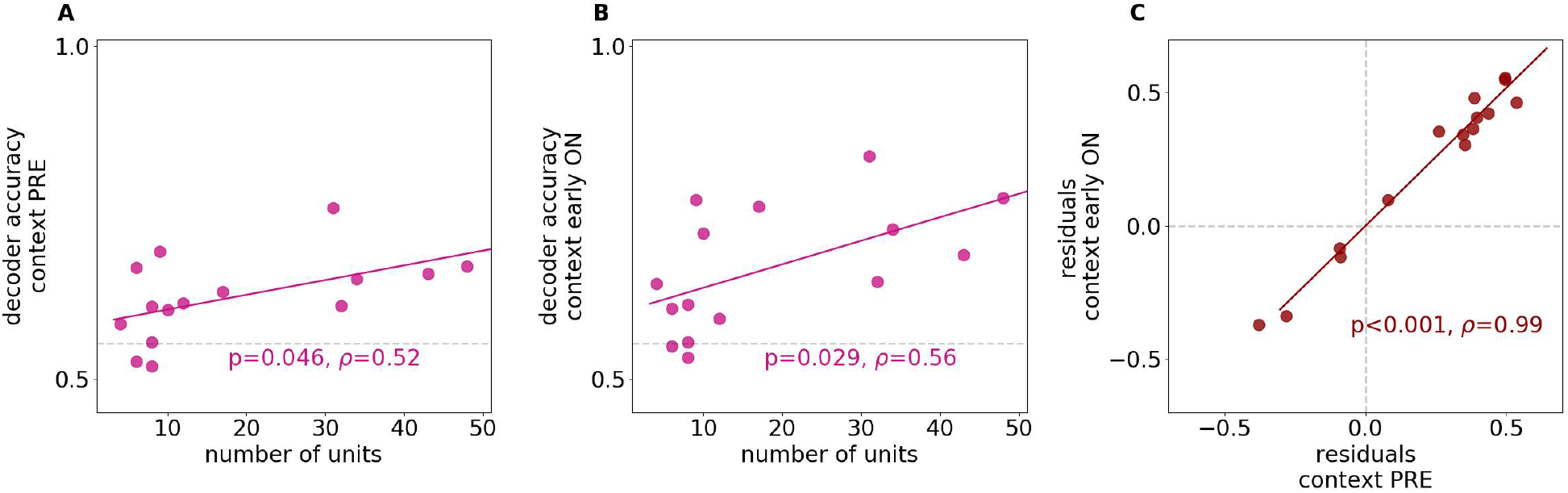
Control for number of neurons in context representation. **A**, Accuracy of a decoder cross-validation test accuracy for a task variable is predicted from number of units. Dots represent average accuracy of context decoders before stimulus onset for a single animal. Grey lines are animal-averaged randomized chance levels as previously. **B**, same as A, but regressed for context average accuracy during stimulus. Grey lines are animal-averaged randomized chance levels as previously. **C**, residuals from the fits from A and B are correlated, containing no relation between number of neurons and accuracies.

**Supplementary Figure 5.**
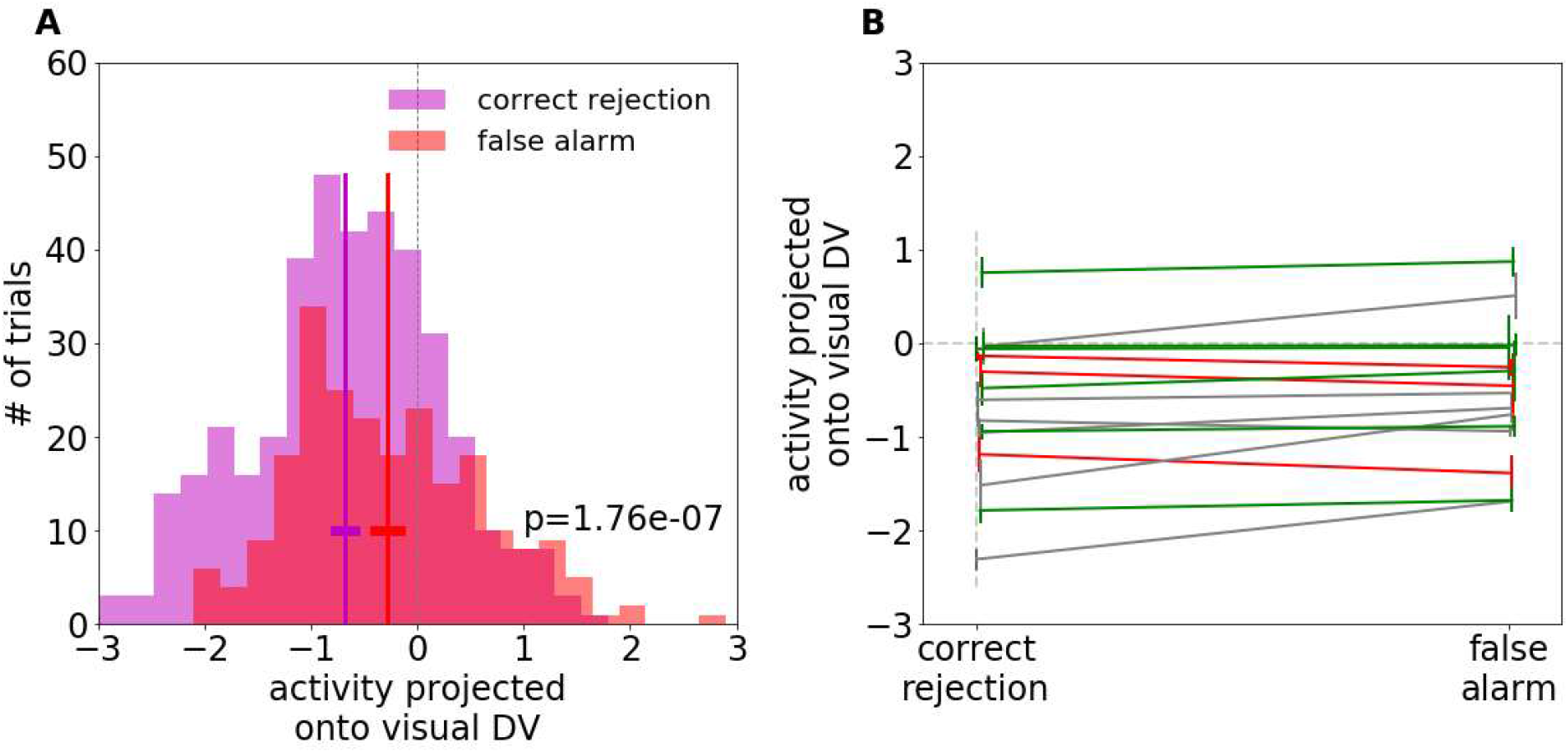
Visual discrimination in error trials. **A**, Histogram of neural activity projected onto standardized visual DVs in no-go attend visual trials, aggregate from all mice. Purple and red colors represent correct and erroneous choices respectively. Vertical lines: mean, horizontal lines 2 s.e.m. Two-sampled t-test for non-equality of means is highly significant. **B**, Means for correct and error trials from **A**, but calculated for each animal separately (lines), caps show 2 s.e.m. Colors show statistics for visual discrimination by neural activity projections found by behaviour conditioned visual decoders: significantly worse discrimination (green), better discrimination (red) in error trials compared to correct trials, or overlapping means (grey). Horizontal point coordinates slightly randomized for ease of visibility.

